# Molecular Dynamics Simulations of Ion Permeation in human NaV Channels

**DOI:** 10.1101/2022.09.29.509656

**Authors:** Giulio Alberini, S. Alexis Paz, Beatrice Corradi, Cameron F. Abrams, Fabio Benfenati, Luca Maragliano

## Abstract

The recent determination of cryo-EM structures of voltage-gated sodium (Na_v_) channels has revealed many details of these proteins. However, knowledge of ionic permeation through the Na_v_ pore remains limited. In this work, we performed atomistic molecular dynamics simulations to study the structural features of various neuronal Na_v_ channels based on homology modeling of the cryo-EM structure of the human Na_v_1.4 channel and, in addition, on the more recent resolved configuration for Na_v_1.2. In particular, single Na^+^ permeation events during standard MD runs suggest that the ion resides in the inner part of the Na_v_ selectivity filter (SF). On-the-fly free-energy parametrization (OTFP) temperature accelerated molecular dynamics (TAMD) was also used to calculate two-dimensional free energy surfaces (FESs) related to single/double Na^+^ translocation through the SF of the Na_v_1.2 homology model. The same thermodynamic analysis was performed on the cryo-EM based Na_v_1.2 configuration. These additional simulations revealed distinct mechanisms for single and double Na^+^ permeation through the wild-type SF, which has a charged lysine in the DEKA ring. In particular, the extracted protein-ion configurations are not accessible by the other modified SFs tested by TAMD/OTFP. Overall, the description of these mechanisms gives us new insights into ion conduction in human Na_v_ cryo-EM based configurations, that could advance understanding of these systems and how they differ from potassium and bacterial Na_v_ channels.

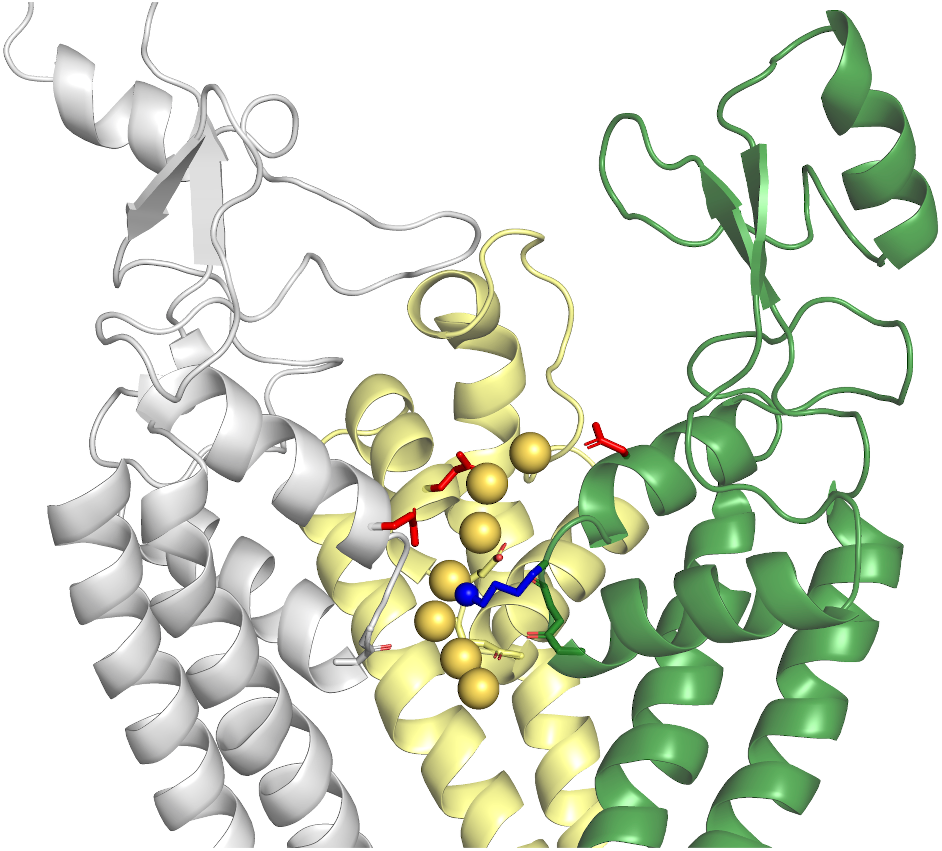

## Introduction

Voltage-gated sodium (Na_v_) channels are integral proteins that are responsible for Na^+^ conduction across the membrane of various excitable cells, where they participate in essential functions including the triggering of neuronal action potential.^1–7^ Several Na_v_ mutations induce dysfunctions linked to a variety of pathological conditions such as epileptic seizures, motor neuron disease and pain,^8–13^ making them a major target for the design of efficient therapeutic strategies.^14–19^ Despite their physiological importance, little is known on their structural mechanisms, also because of the limited availability of human Na_v_ structures. The Na_v_ channels family includes both prokaryotic (bacterial) and eukaryotic members, which are evolutionary divergent even if they share the same main main functions.^20,21^ In particular, the eukaryotic sub-family of human Na_v_ proteins comprises nine homologs, named Na_v_1.1 to Na_v_1.9, all characterized by highly conserved sequences.^22–24^ These channels show tissue specific distribution, with Na_v_1.1, Na_v_1.2 and Na_v_1.6 expressed in the central nervous system (CNS).^15^ Moreover, unlike bacterial Na_v_ channels, made up of four separated subunits, the human Na_v_ channels are formed by four different domains, also called repeats (indicated as Rep. DI-DIV), connected together to form a single long polypeptide chain known as the ion-conducting *α*-subunit. When viewed from the extracellular side, the DI-DIV domains are assembled in a clockwise fashion, each containing six transmembrane (TM) *α*-helices, named S1 to S6. The S1-S4 segments of each repeat form four independent voltage sensor domain (VSDs),^5,25–27^ while the last two S5-S6 helices are assembled to generate the pore domain (PD).^28–38^ Between the S5 and S6 segments there is a motif (named P-loop) that includes the Na^+^ selectivity filter (SF),^15,39^ which ensures the selective permeation of Na^+^ ions and differs between bacterial and eukaryotic channels. In particular, bacterial Na_v_ channels display a ring of four E side chains (EEEE),^39^ as evidenced by high-resolution crystallographic structures. ^40–44^

Several independent studies used molecular dynamics (MD) simulations to investigate the structural properties of bacterial Na_v_ proteins.^45^ According to Ref.s 34,46, the main results achieved could be summarized as follows.

- A first group of simulations revealed information regarding the binding sites in the SF and their multi-ion occupancy.^47–60^
- Additional microsecond time-scale MD simulations showed that the SF symmetric configuration of the crystal structures can be broken, affecting both the permeation and the selectivity mechanisms. ^53,58,59,61,62^

The number of Na^+^ ions that permeate the SF is also a central issue. The observation of a multi-ion process is in agreement with the recent crystallographic structures of a bacterial Na_v_Ms channel introduced in Ref. 44. Moreover, the X-ray crystal structure^42^ of the closed conformation of Na_v_Ae1p, another prokaryotic Na_v_ channel, is characterized by an outer ion site in the SF that suggests multiple ion-binding sites. ^15^

Mammalian Na_v_ channels replace the EEEE motif with a D_I_E_II_K_III_A_IV_ ring^39,63–65^ (here, following the notation in Ref. 46, subscripts indicate the Na_v_ domain associated to each residue) including a lysine side chain which selectively slows the transport of K^+^ over Na^+^. ^66–69^ The SF is formed by the four side chains of the D_I_E_II_K_III_A_IV_ signature at the extracellular side, and, at the inner side, by the couple of backbone carbonyl oxygen atoms belonging to the two preceding residues in each repeat. ^63–65^ Indeed, since they are not involved in hydrogen bonds with other residues, these two inner rings of carbonyls are free to interact with the permeating cations.^63–65,70^ In addition, both experimental results and Monte Carlo simulation studies suggested the pivotal role of the E and K residues in the DEKA ring. ^66,70^ Several investigations demonstrated that the specific position of the residues in the SF is essential to preserve the correct selectivity. ^66,71^

Of particular interest, the DEEA variant^22,72–77^ reverts the SF to an ancestral, Ca^2+^selective, state. In the case of the Na_v_1.2 channel, the alteration of the third residue in the DEKA motif (K1422E) affects the organism with both loss-of-function and gain-of-function effects.^67,71^ In addition to the DEKA signature, human channels (with the exception of Na_v_1.7) display a well-conserved^65,78^ outer ring consisting of four charged residues, E_I_E_II_D_III_D_IV_, (again, subscripts indicate the Na_v_ domain associated to each residue^46^), supposed to increase the electrostatic attraction and conductance of extracellular cations, but not selectivity. ^71,79^ Remarkably, the residues in both the EEDD and DEKA motifs are asymmetric in their position along the filter axis and characterized by a high degree of flexibility.^65^ To cope with such structural complexity, in the following we will refer to the region including the conductive EEDD signature, the DEKA motif and the two inner rings as the conductivity/selectivity filter (C/SF). As for ionic occupancy, although early experiments suggested that Na_v_ channels are characterized by multiple binding sites, ^80–83^ simultaneous binding of multiple ions is not universally accepted, leading to disagreements on the mechanism of ionic conduction. ^80–85^ Previous computational studies examined also the role of the lysine belonging to the DEKA ring in conduction and selectivity, but they were based on models using modified bacterial structures^46,86,87^ or homology models of mammalian Na_v_ channels from bacterial templates 88–91. In two separate articles,^86,87^ the authors replaced the ring in the bacterial Na_v_Rh channel with the DEKA motif and performed MD simulations, describing the screening process of the K residue, which pushes the Na^+^ ion toward the D/E residues. On the other hand, the MD studies of Ref.s 88–91, based on an homology model of the rat Na_v_1.4 pore-only channel from the bacterial Na_v_Ab structure, showed an essential displacement of the lysine in order to allow proper ionic permeation. More recently, the results based on extended MD simulations in Ref. 46 suggest that the role of lysine in conduction and selectivity is strongly dependent on its protonation state. When the K residue is deprotonated (*i.e*. uncharged), 2 and 3-ion occupancies are preferred. On the contrary, when the residue is protonated (positively charged), a reduced occupancy (1 and 2) is observed.

Despite these seminal works, the investigation of the ionic permeation mechanisms in Na_v_ channels remains mostly focused on bacterial-based models, and the transferability of these results to the more recently solved experimental Cryo-EM structures of eukaryotic Na_v_ proteins^63–65^ has been investigated only in few works. ^70,92–94^ Therefore, a thermodynamic characterization of the coupling between ionic permeation and the K residue in the DEKA motif is still missing.

In this work, we aim to investigate the residence of Na^+^ ions in structures of neuronal Na^+^ channels by means of MD simulations and free energy (FE) calculations. To this aim, we used both homology-based Na_v_ structures from Cryo-EM templates and experimental CryoEM Na_v_ structures. Starting from the first Cryo-EM structure of a human Na_v_ channel (Na_v_1.4, PDB ID: 6AGF,^65^ at 3.20 Å resolution) we obtained the homology-modeled structures of the *α* subunits of three neural Na_v_ channels (Na_v_1.1, Na_v_1.2, Na_v_1.6). We performed all-atom unbiased MD simulations to assess the structural stability of these structures in the wild type (WT) configuration immersed in a water-membrane environment at a 150 mM physiological concentration of NaCl. Na^+^ permeation events from the external space to the internal cavity have been observed in all the systems. Then, focusing on the Na_v_1.2 model, as a paradigmatic example of neurogical Na_v_s, we used pore-only configurations (*i.e*., without the VSDs) to study the Na^+^ permeation in the C/SF via FE surface (FES) calculations using the temperature accelerated MD/on-the-fly free-energy parametrization (TAMD/OTFP) method. ^95–98^ Special attention was devoted to the role of SF motif in the WT and DEEA structure, considering both the charged and uncharged version of the K or E1422 residue. The investigation was replicated on the more recent Na_v_1.2 Cryo-EM structure (PDB ID: 6J8E,^99^ at 3.00 Å resolution, based on the aforementioned Na_v_1.4 structure). The overall conformation of the experimental Na_v_1.2 is substantially identical to the previous Na_v_1.4 template with a root-mean-square deviation (RMSD) of 0.696 Å over 981 Cα atoms, according to the analysis exposed in Ref. 99. Based on this observation, not surprisingly, the results of the MD simulations performed with the latter structure nicely fit with those obtained with the starting homology-based model. Our simulations describe distinct mechanisms for single and double Na^+^ ions during permeation through the WT filter, with the charged K1422 residue. Remarkably, in the case of the WT configuration, both standard MD runs and FE calculations highlight single Na^+^ permeation events, in the course of which the ion stays in a global FE minimum in the inner part of the Na_v_ SF. These configurations observed for the WT with the charged K1422 residue are not accessible to all the other simulated systems. Our results provide a first attempt to describe the permeation mechanisms of Na^+^ translocation through neuronal human Na_v_ channels, using the information provided by the recent Cryo-EM human Na_v_ structures. Globally, these outcomes could help in describing the complex mechanism of ion permeation of these systems.

## Materials and Methods

### Structural modeling and standard MD simulations

#### Structural modeling - template choice

When we started this project, no experimental configuration was available for mammalian neuronal Na_v_1.1, 1.2 and 1.6 channels. Therefore, we first relied on homology-based modeling to build accurate structural models for these proteins. The cryo-EM structure of the human Na_v_1.4 *α*-protein, in complex with the auxiliary *β*1 subunit, introduced in Ref. 65 (PDB ID: 6AGF), has been used as template. This structure is characterized by a detailed description of the pore domain, including the C/SF that regulates the permeation of ions. In particular, the critical DEKA motif is located in the inner part of the cavity and was reliably resolved with a local resolution of up to ~ 2.8 Å. In **Table 1**, we introduce the sequence of the different domains modeled for each channel. The structures include all the four Rep. DI-DIV TM domains, plus the intracellular link between DIII and DIV (III-IV link). The long extracellular loop located in the DI part was not included in the modeling because unresolved in the template. Therefore, the residues adjacent to the missing region were connected forming one single DI polypeptide chain.

**Table 1:** List of all the residues included in homology modeling. The starting UNIPROT ID with the associated human sequence is included.

| Channel | Na<sub>v</sub>1.1 | Na<sub>v</sub>1.2 | Na<sub>v</sub>1.6 |
| --- | --- | --- | --- |
| UNIPROT ID | P35498 | Q99250 | Q9UQD0 |
| Rep. 1 | 129-420 (no 285-305) | 125-427 (no 285-305) | 133-408 (no 290-299) |
| Rep. 2 | 769-991 | 755-983 | 754-976 |
| Rep. 3 | 1220-1479 | 1210-1473 | 1200-1459 |
| III-IV Link | 1480-1542 | 1474-1526 | 1460-1523 |
| Rep. 4 | 1543 -1785 | 1527 -1776 | 1524-1765 |

#### Notation of the SF configurations

Here we report the notation used to differentiate the Na_v_ SF configurations investigated in this work:

- **DEK**^+^**A**, WT configuration of the DEKA ring, with the positively charged lysine.
- **DEK**^0^**A**, WT configuration of the DEKA ring, with uncharged lysine.
- **DEE**^−^**A**, mutated configuration of the DEKA ring, with the charged glutamate in place of the lysine.
- **DEE**^0^**A**, mutated configuration of the DEKA ring, with uncharged glutamate in place of the lysine.

#### Structural modeling of neuronal Na_v_ channels

The *α*-subunit of human Na_v_ channel are formed by four TM domains linked by cytoplasmic loops. Our TM structural models of human neuronal Na_v_ channels were constructed in two steps^100^ following a procedure similar to the one introduced in Ref.s 101,102. Firstly, we modeled each of the four TM DM domains separately, using the advanced protein prediction method included in the SWISS MODEL server^103^ with the exemption of the Rep. III and Rep IV. These domains were modeled together, including also the cytosolic III-IV Link. The conservation between the sequence of the template and the domains in each of the three neural Na_v_ channels spans from ~ 85 % to ~ 91 %. Then, at a second stage, the whole subunit for each channel was built by structurally aligning four individuals to the template in a clockwise order viewed from extracellular side, using UCSF Chimera.^104^ The resultant, tetrameric model was refined again by FG-MD^105^ to eliminate unwanted inter-domain steric clashes and improve the global model quality. Before MD simulations, all the systems were aligned to the membrane normal axis *z*, using the OPM-PPM^106^ service available at opm.phar.umich.edu/ppm_server. We show the homology-based Na_v_1.2 model in **Figure 1**, including the representation of the asymmetric Na_v_1.2 SF. The Na_v_1.1 and Na_v_1.6 SFs are also introduced in **Figure S1** and **Figure S2**, respectively.

**Figure 1:**
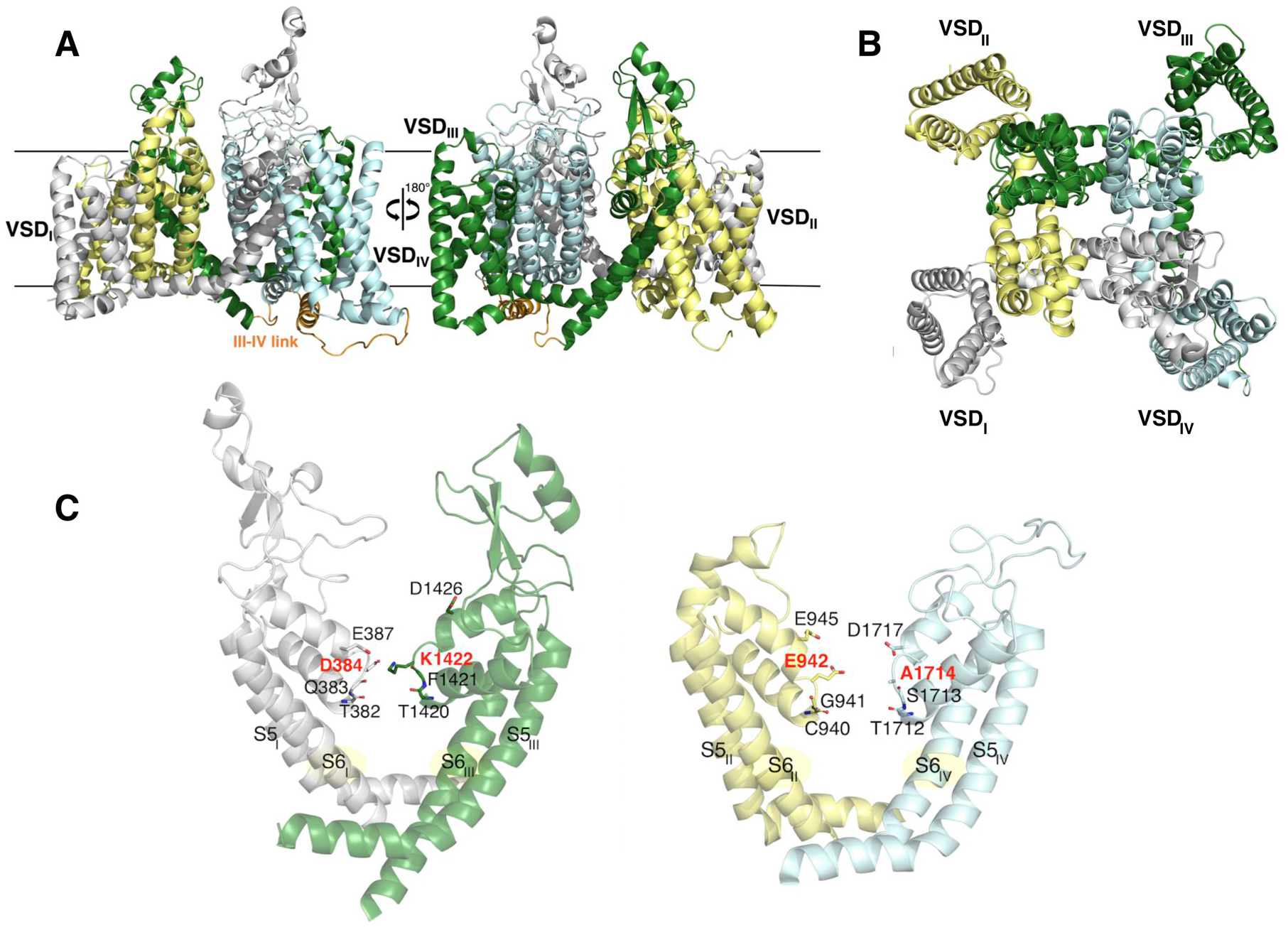
**A.** The equilibrated homology-based Na_v_1.2 model. The domains are colored as in Ref.s 63–65: Domain I, gray; Domain II, yellow; Domain III, green; Domain IV, cyan. This color scheme is applied throughout the manuscript. **B.** Extracellular view of the model. **C.** Representation of the asymmetric Na_v_1.2 SF. The SF vestibule is enclosed by the side chains of the D_I_E_II_K_III_A_IV_ motif (D384/E942/K1422/A1714) and the internal carbonyl oxygen atoms of the two preceding residues in each repeat. The residues of the external E_I_E_II_D_III_D_IV_ motif (E387, E945, D1426, D1717), above the DEKA domain, are also shown. The EEDD domain and the SF region form the C/SF.

#### MD of the WT Na_v_1.2 model - standard atomic masses set up

We performed a 500 ns long MD simulation of the human Na_v_1.2 (hNa_v_1.2) structural model, to assess the quality of the modeling protocol. CHARMM hydrogen atoms were added with CHARMM-GUI and disulfide bonds were assigned according to the information provided in the PDB structures 5XSY, 6AGF and 6J8E. In particular, the disulfide bond between C950 and C959, which is located inside the pore cavity, was included in all the simulations of this work. The SF filter was modeled with a DEK^+^A ring. Then, the Na_v_1.2 structure was inserted in a homogeneous lipid patch of 1-palmitoyl-2-oleoyl-sn-glycero-3-phosphocholine (POPC) membrane environment with explicit water molecules and the total charge was neutralized with a 150 mM NaCl solution, obtaining ~ 290, 000 atoms in total. For this MD simulation, as for the other standard MD runs described in this work, we used the NAMD software^107^ (version 2.12) and the CHARMM36m^108–110^/CHARMM36^111^ parameters for the protein and lipids, respectively, together with TIP3P model for water molecules^112^ and the associated ionic parameters with NBFIX corrections.^113–115^ Tetragonal Periodic Boundary Conditions (PBCs) were applied to the simulation box to remove surface effects. Long range electrostatic interactions were calculated using the Particle Mesh Ewald (PME) algorithm.^116^ Electrostatic and van der Waals interactions were calculated with a cutoff of 12 Å and the application of a smoothing decay starting to take effect at 10 Å. A time-step of Δ*t* = 2 fs was employed. To ensure maximum accuracy, electrostatic and van der Waals interactions were computed at each simulation step. All covalent bonds involving hydrogen atoms were constrained using the SHAKE/SETTLE algorithms.^117,118^ Before production, the system was relaxed following a ~ 35 ns equilibration, by extending the CHARMM-GUI equilibration protocol, in order to allow proper hydration of solvent exposed regions of the Na_v_ pore cavity. Then, the system was simulated in the NPT ensemble using the Nosé-Hoover Langevin piston method^119,120^ to maintain the pressure at 1 atm, and a Langevin thermostat at 310 K. The oscillation period of the piston was set at 50 fs, and the damping time scale at 25 fs. The Langevin thermostat was set with a damping coefficient of 1 ps^−1^.

#### MD of the WT Na_v_1.1, Na_v_1.2, Na_v_1.6 models - the HMR set up

The three neuronal models Na_v_ 1.1, 1.2 and 1.6 were simulated in the WT configuration. In all these standard simulations, the K residue (K1422 in the Na_v_ 1.2 channel) was studied in the charged configuration (DEK^+^A) and the hydrogen mass repartitioning (HMR) method was used to build the topology of the system. HMR is a computational methodology that re-distributes the atomic masses.^121,122^ Thanks to this scaling, a larger time-step can be used in the simulations. The method was originally proposed in Ref. 121 and then successfully applied to protein simulations in Ref. 122, using a hydrogen mass of 3 amu and a Δ*t* = 4 fs. For our simulations, CHARMM hydrogen atoms were added with CHARMM-GUI and disulfide bonds were assigned as done for the previous system, including the disulfide bond between C950 and C959 in Na_v_1.2 (C959-C968 in Na_v_1.1 and C944-C953 in Na_v_1.6, respectively), which is located inside the pore cavity. Then, the Na_v_ structures were inserted in a POPC membrane environment and solvated with explicit water molecules and a 150 mM NaCl solution. For all these MD simulations, we used the NAMD software^107^ and the parameters of the previous simulation with standard atomic masses. Tetragonal Periodic Boundary Conditions (PBCs) were applied to the simulation box to remove surface effects. Long range electrostatic interactions were calculated using the Particle Mesh Ewald (PME) algorithm.^116^ Electrostatic and van der Waals interactions were calculated with the standard CHARMM cutoff of 12 Å and the application of a smoothing decay starting to take effect at 10 Å following the instructions in Ref. 123. A time-step of 4 fs was employed coupled to the HMR scheme. Electrostatic and van der Waals interactions were computed at each simulation step. All covalent bonds involving hydrogen atoms were contrained using the SHAKE/SETTLE algorithms.^117,118^ Before the production, the systems were relaxed following a ~ 35 ns equilibration. Then, the systems were simulated in the NPT ensemble using the Nosé-Hoover Langevin piston method^119,120^ to maintain the pressure at 1 atm, and a Langevin thermostat at 310 K. The oscillation period of the piston was set at 300 fs, and the damping time scale at 150 fs. ^123^ The Langevin thermostat was employed with a damping coefficient of 1 ps^−1^.

#### MD of the WT Na_v_1.2 Cryo configuration - the HMR set up

We used also an experimental Cryo-EM structure of the human Na_v_1.2 channel. The structure is characterized to be bound to the peptidic *μ*-conotoxin KIIIA pore blocker (PDB ID: 6J8E).^99^ Notably, the structure includes a Na^+^ ion in the C/SF, as illustrated in **Figure 2**. Moreover, the bottom tip of the KIIIA peptide includes only one residue, K7, whose amine group interacts with E945.^99^ The overall conformation of the experimental-based Na_v_1.2 is substantially identical to the Na_v_1.4 template with a root-mean-square deviation (RMSD) of 0.696 Å over 981 C*α* atoms, according to the analysis exposed in Ref. 99. Based on this observation, this structure nicely fits with our homology-based Na_v_1.2 model. The superposition of the two structures is illustrated in **Figure S3**. The comparison between the Cryo-EM based Na_v_1.2 and our preliminary homology-based model is characterized by an RMSD of 0.746 A over the Ca atoms belonging to the transmembrane *α* helices. Furthermore, the pore only configuration of the new Na_v_1.2 structure differs from our pore-only model by an RMSD of 0.534 Å. To further investigate the structural features of this new configuration, we produced two MD runs of the *α*-subunit of this structure (after removing all the non *α*-subunit atoms, including the toxin and the ion). In particular, we performed 2 × 500 ns MD simulations with a physiological 150 mM NaCl concentration, using the HMR protocol illustrated in the previous paragraph.

**Figure 2:**
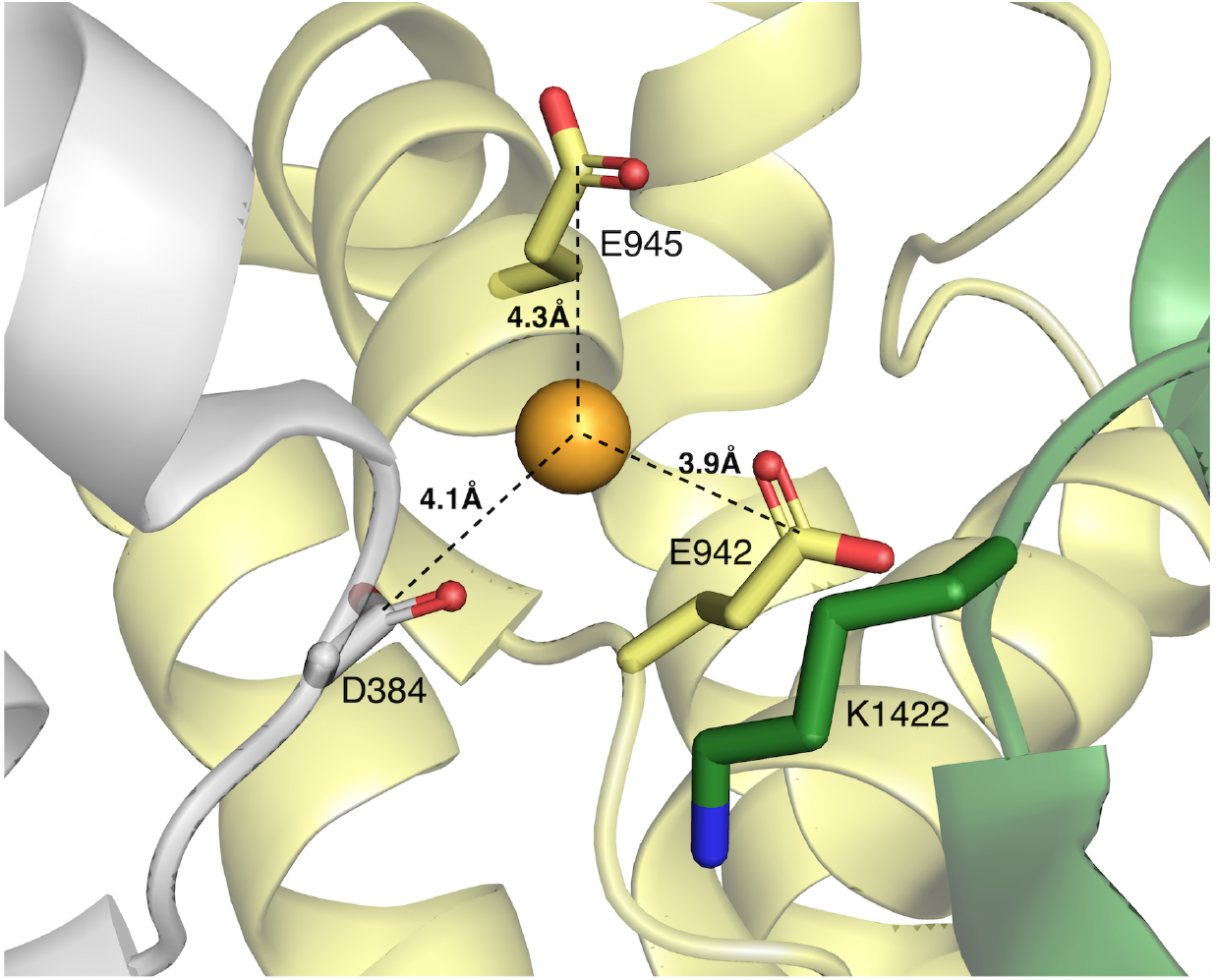
Sodium ion resolved in the SF of the PDB 6J8E Na_v_1.2 Cryo-EM structure. The pore blocker *μ*-conotoxin KIIIA is not shown.

#### Ion permeation events

For each standard MD simulation listed in **Table S1**, we monitored the ionic permeation events through the SF in the presence of a physiological concentration of 150 mM of NaCl. The calculation includes the volume of the C/SF. Here we report the details of the selections using the notation of the Na_v_1.2 residues. Similar selections were made for the Na_v_1.1 and Na_v_1.6 based systems. The selected domain is characterized by the following features:

- a radius of 7 Å around the pore axis;
- the quota of the upper circular section is defined by the *z* coordinate of the C*α* atom following the E939 residue of the EEDD domain;
- the quota of the lower circular section is defined by the *z* coordinate of the C*α* atom of residue 940, further reduced by 4 Å.

#### Distributions of atomic positions

We analyzed the positions within the C/SF of one sodium ion, and the lysine sidechain of the DEKA motif. To this goal, we combined all six MD runs of the WT systems in NaCl solution, for a total of 3 *μ*s. For each dataset, we reconstructed the histogram of values of the *z*-coordinate of a single Na^+^, provided that they were found inside the C/SF. The same analysis was performed for the DEKA lysine’s NZ atom, which is supposed to play a role in ionic permeation through mammalian Na_v_ channels.^46,86–91^ The aggregation of the results from various systems is justified by the fact that the homology based Na_v_1.2 and the Cryo-EM based Na_v_1.2 structure are nearly identical in the C/SF and, because this is a highly conserved region in the various Na_v_ proteins, the identity is transferred also to the other channels we built. Therefore, in terms of this analysis, we considered the various neuronal Na_v_ channels as equivalent systems.

### TAMD/OTFP simulations for free energy calculations

The free energy (FE) of an *N*-atom system, with potential energy *U*(***x***) and at temperature *T*, is defined as

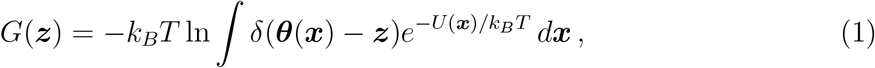

where the integral runs over the system configurations 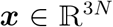, and 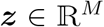 represents a point in the lower-dimensional space of CV obtained via the transformation 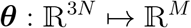, with *M* < 3*N*. In temperature-accelerated molecular dynamics (TAMD),^95^ a set of *M* auxiliary variables ***z*** = *z*_1_,…, *z_M_* are introduced and tethered to the *M* collective variables (CVs) ***θ*** = *θ*_1_,…, *θ_M_*. The combined set (***x***, ***z***) is subjected to the following potential:

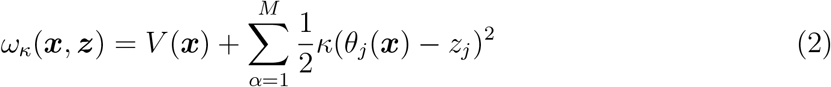

where *κ* > 0 is an adjustable spring-constant-like parameter. The extended system evolves according to coupled equations, as for example:

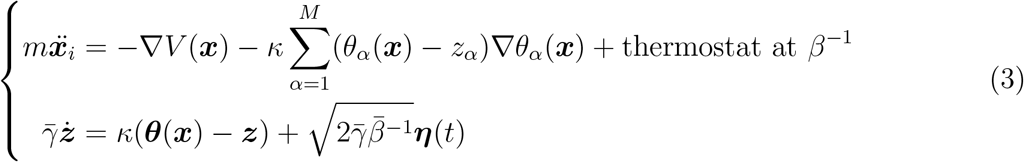

where *m* is the mass, ***η***(*t*) is a Gaussian process with mean 0 and covariance 〈*η_α_*(*t*)*η_α′_*(*t*′)〉 = *δ_αα′_δ*(*t* – *t*′), 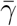 is a friction coefficient and 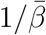 is an artificial temperature with 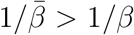, where 1/*β* is the inverse of the temperature *k_B_T*. As shown in Ref. 95 (see also Ref. 124), by adjusting the parameter *κ* so that ***z*** ~ ***θ***(***x***(*t*)) and the friction coefficient 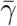 so that ***z*** moves slower than ***x***, a trajectory ***z***(*t*) can be obtained which moves at the artificial temperature 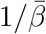 on the FE surface (FES) calculated at the physical 1/*β*. Indeed, it has been demonstrated^95^ that under such assumptions each auxiliary variable *z_j_* is driven by the force

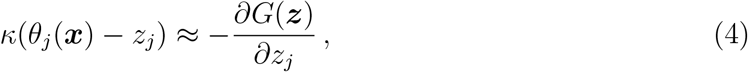

where *G*(***z***) is given by Eq. 1. Hence, by choosing 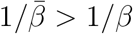 in Eq. 3, the ***z*** trajectory will rapidly visit the regions where the FE is relatively low, overcoming barriers that the system would take a long time to cross at the physical temperature 1/*β*. In addition, the on-the-fly free-energy parametrization (OTFP) method is an efficient approach to reconstruct the FES sampled by a TAMD simulation.^96–98^ The idea is to introduce an approximation of the FE via a set of basis functions *ϕ_m_*(***z***):

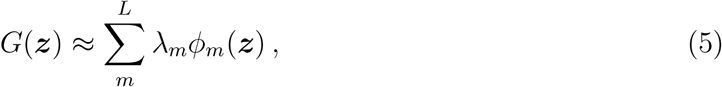

and to determine the coefficients in Eq. (5) by minimizing the following error function^125^

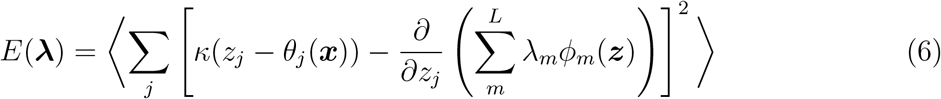

where λ is the 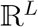 vector of the λ_*m*_ coefficients and 〈〉 indicates the average computed during the TAMD simulation. This is equivalent to solve a linear system of equations, ***A*λ** = ***b*** where

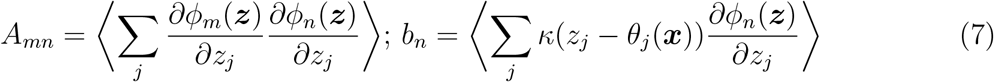

Following Ref.s 96–98,126, chapeau functions are used as basis set. Other FE reconstruction methods based on the minimization of the same error function have been introduced more recently.^127,128^

The NAMD code^107,129^ was used to integrate the atomic equations of motion in the OTFP simulations, while the auxiliary equations and the reconstruction is handled by a C-code via the NAMD TcL forces plugin,^98,130^ available at github.com/cameronabrams/otfp. The reduced pore-only configuration of the former Na_v_1.2 model used for the TAMD/OTFP calculations includes the following residues: Rep. 1: from 234 to 427, with the exclusion of residues from 285 to 305; Rep. 2: from 866 to 983; Rep. 3: from 1320 to 1476; Rep. 4: from 1641 to 1775. The protein was aligned to the membrane normal *z* axis, using OPM-PPM.^106^ The same pore configuration was obtained from the Cryo-EM Na_v_1.2 structure (PDB ID: 6J8E). In order to use absolute coordinates as CVs for both systems, they were aligned by superimposing the Cryo-EM pore on the model one. As reference, the *z*-coordinate of the Cryo-EM K1422 NZ atom after alignment is 6.8 Å. The resolved Na^+^ ion in the SF (at *z* = 12.7 Å) was removed from the PDB file. Then, both systems were inserted in a homogeneous POPC membrane, solvated with explicit water molecules and the total charge was neutralized with a 150 mM NaCl solution. The minimum starting size of the periodic box measured ~ [135 × 135 × 129] Å^3^, which ensured a distance larger than 20 Å between adjacent images of the protein during the simulations. For all the TAMD/OTFP simulations, we employed the CHARMM parameters already used for the standard MD simulation with original masses. Tetragonal Periodic Boundary Conditions (PBCs) were applied to the simulation box and long range electrostatic interactions were calculated using the Particle Mesh Ewald (PME) algorithm. ^116^ Electrostatic and van der Waals interactions were calculated with a cutoff of 12 Å and the application of a smoothing decay starting to take effect at 10 Å. A time-step of 2 fs combined with the SHAKE/SETTLE algorithms^117,118^ was used. In order to ensure maximum accuracy electrostatic and van der Waals interactions were computed at each simulation step. Before production, the system was equilibrated for ~ 25 ns. A snapshot of the equilibrated system is illustrated in **Figure 3**, panel **B**. During all the TAMD/OTFP runs, soft harmonic restraints were maintained on the C*α* atoms of each *α* S5-S6 helix in order to avoid any rigid body rotational or translational displacement of the protein. More specifically residues from 234 to 271 and from 399 to 427 in Rep. DI; residues from 866 to 907 and from 956 to 983 in Rep. DII; residues from 1321 to 1359 and from 1447 to 1476 in Rep. DIII and residues from 1643 to 1682 and from 1749 to 1775 in Rep. DIV were restrained. On the other hand the C/SF was completely free to allow proper sampling. After the preparation, the systems were simulated in the NPT ensemble using the Nosé-Hoover Langevin piston method^119,120^ to maintain the pressure at 1 atm, and a Langevin thermostat at 310 K. The oscillation period of the piston was set at 50 fs, and the damping time scale at 25 fs. The Langevin thermostat was employed with a damping coefficient of 1 ps^−1^. The ionization state of the protein residues correspond to the neutral pH (DEK^+1^A and DEE^−1^A) except for the simulations that include the modified protonation of the K/E1422 residue (DEK^0^ and DEE^0^A). All the runs where performed in a zero-transmembrane voltage regime. All the cations not involved in the definition of the CVs were excluded from the C/SF by applying a repulsive restraint. This restraint applies in a rectangle positioned approximately at the center of the DEK(E)A motif by including the following bias to the force field expression of the potential energy:

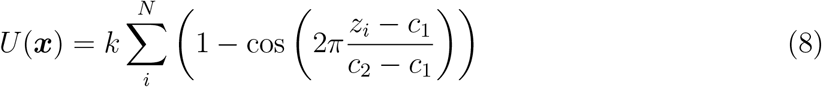

where the summation runs over the excluded cations inside the SF, i.e., inside the rectangle delimited by the following coordinates {*x_min_*, *y_min_*, *z_min_*} = {-6 Å, −6 Å, −4 Å}, and {*x_max_*, *y_max_*, *z_max_*} = {6 Å, 6 Å, 18 Å}, which was used for the majority of the simulations, as reported in **Table S2**. Here, *c*_1_ = −4 Å, *c*_2_ = 18 Å, and *k* = 600 kcal/mol. This potential introduces a prohibitive barrier for the excluded molecules to enter the SF. The same potential term has been previously used in Ref. 98 to perform TAMD/OTFP simulations of a representative voltage gated potassium K**v** channel while keeping water molecules outside the K_v_ SF. The single or two ions used for defining the CV were confined inside the filter by the application of two 300 kcal/mol half-harmonic potentials at *z* > 17 Å and *z* < 4 Å.

**Figure 3:**
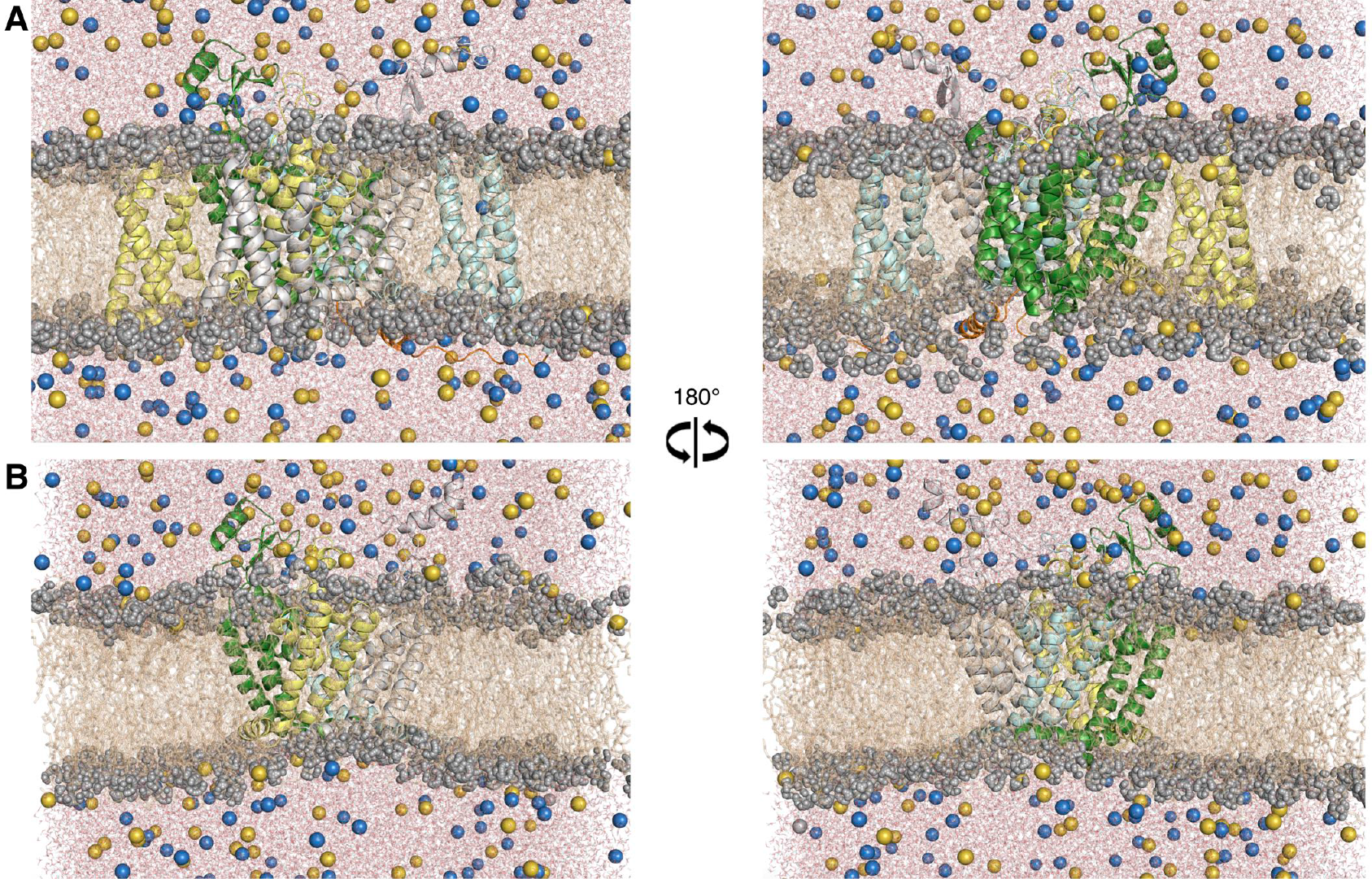
**A.** The equilibrated homology-based Na_v_1.2 *α* subunit embedded in a hydrated POPC bilayer (brown wires), surrounded by water (red and white). Sodium (yellow) and Chloride (blue) ions are represented as spheres. **B.** The equilibrated homology-based Na_v_1.2 pore-only domain embedded in the same environment described in the previous panel.

Two dimensional free energy surfaces (2D-FESs) were recovered by combining the following CVs:

- *z*_1_: the *z*-coordinate of one Na^+^ in the C/SF. This CV will be denoted as *r_z_*(Na_1_);
- *z*_12_: the *z*-coordinate of the center of mass (COM) of two Na^+^ ions in the C/SF. This CV will be denoted as COM_*z*_(Na_1_, Na_2_);
- *z_NZ_*: the *z*-coordinate of the NZ atom of the K1422 sidechain. This CV will be denoted as 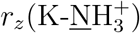 in the DEK^+^A and *r_z_*(K-<u>N</u>H_2_) in the DEK^0^A, respectively;
- *z_CD_*: the *z*-coordinate of the COM of the CD atom of the E1422 sidechain. This CV will be denoted as *r_z_*(E-<u>C</u>OO^−^) in the DEE^−^A and *r_z_*(E-<u>C</u>OOH) in the DEE^0^A, respectively.

In the following, we will describe two sets of TAMD/OTFP simulations. The first, denoted with OTFP_1_, considers *z*_1_ and either *z_NZ_* or *z_CD_* depending on the composition of the 1422 residue. The second one, denoted with OTFP_2_, considers *z*_12_ and either *z_NZ_* or *z_CD_*. The auxiliary temperature was 6000 K, the auxiliary friction 800 (kcal/mol) ps/Å^2^, and the spring constant 1500 kcal/mol Å^−2^. The OTFP_1_ and OTFP_2_ set up were produced for all the SF configurations included in this work (DEK^+^A, DEK^0^A, DEE^−^A and DEE^0^A), in the presence of a 150 mM NaCl concentration, using both the Na_v_1.2 model and the Cryo-EM based Na_v_ 1.2 structure. A list of the OTFP simulations is reported in **Table S2**.

#### Structural analysis and atomic density calculations

All the MD trajectories were visualized and analyzed using UCSF Chimera^104^ (www.cgl.ucsf.edu/chimera/), Pymol^131^ (pymol.org/2/), the NAMD-COLVAR module^132^ and VMD^133^ (www.ks.uiuc.edu/Research/vmd/) with in-house Tcl scripts. The VMD QuickSurf representation^134–136^ was used to represent the volume occupancy of selected atoms in the metastable conformations extracted from the TAMD/OTFP simulations. This VMD plugin computes an isosurface extracted from the expansion of a Gaussian density kernel map at each particle position and uses a 3D grid to sum all together and get the final density. Volume densities were colored according to charged atoms from DEKA and EEDD residues (red for all the negatively charged residues). The density of the K/E1422 residue is represented according to the specific system (blue for DEK^+1^A, cyan for DEK^0^A, red for DEE^−^A, orange for DEE^0^A). Na^+^ density is also included (blue). An equilibrated snapshot of the full system and of the pore only configuration based on the homology Na_v_1.2 model are introduced in **Figure 3**.

## Results and Discussion

This section is organized in two parts. In the first one, we report the results obtained from the standard MD simulations, both in terms of assessment of the quality of the different structures and of the evaluation of the observed Na^+^ permeation events trough the SF. In the second subsection, we describe the thermodynamic features of Na^+^ translocation into the Na_v_1.2 SF, by analyzing the 2D-FESs extracted from the TAMD/OTFP simulations outlined in Materials and Methods. The list of all the simulations included in this work is reported in **Figure 4** and in **Tables S1** and **S2**.

**Figure 4:**
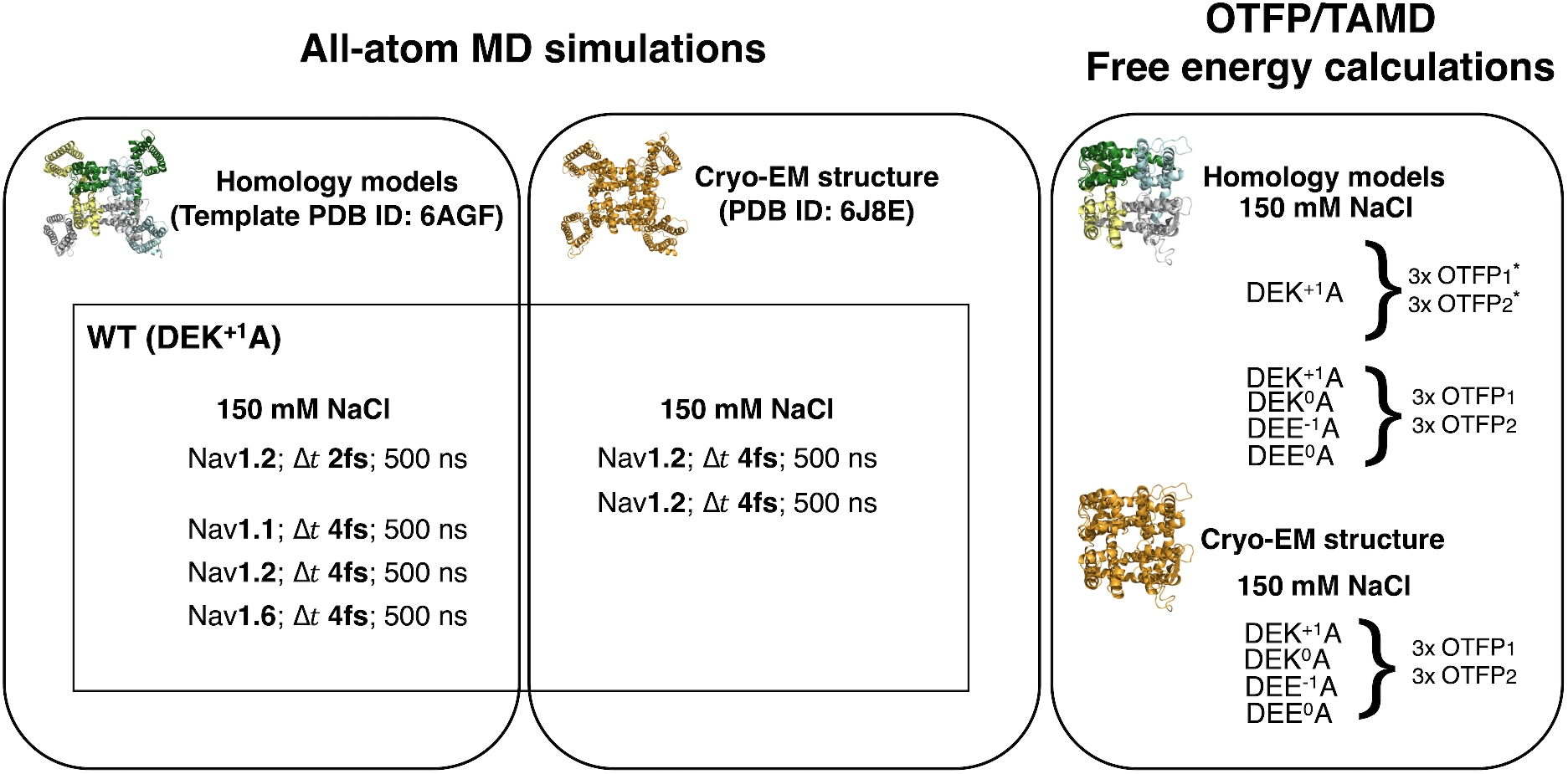
Panel including all the simulations performed in this work. For each MD runs the integration timestep Δ*t* is specified. The meaning of OTFP_1_ and OTFP_2_ is specified in the text.

### Standard MD simulations of the Na_v_ systems

#### Structural stability of the neuronal Na_v_ models and the Na_v_1.2 Cryo-EM structure

We tested the stability of all the Na_v_ structures involved in this study via unbiased MD simulations. For each system, we calculated the backbone RMSD of the transmembrane pore domain, including the S5-S6 helices of each repeat (**Figure S4**). All the RMSD profiles show plateaus at values ~ 2 or 3 Å, thus preserving the overall pore conformation. Remarkably, we found no significant difference between the homology-based and Cryo-EM-based structures of the Na_v_ 1.2 channel. We also calculated a set of cross distances (CDs), each defined between two conserved atoms or groups of atoms belonging to opposite repeats and listed in the Supplementary file. The average values of all CDs, obtained from their mean from each MD run, are reported in **Table S3**. Results for the homology-based models and the Cryo-EM structure are similar.

#### Interactions between sodium ions and the C/SF

Despite the limited sampling of our unbiased MD simulations, they can still provide a partial picture of Na^+^ translocation through the filter. We thus analyzed ion occupancy in the C/SF region, as defined in Materials and Methods. In all the standard MD simulations, we observed various ion permeation events that produced the ionic occupancies reported in **Table 2**. While all Na_v_ systems prefer single Na^+^ occupancy, they also show a small percentage of 2-Na^+^ configurations, as further confirmed by the results exposed in **Table 3**, where we report the average Na^+^ occupancy during each standard MD run.

**Table 2:** Summary of the ionic occupancy (from 0 to 4) in the C/SF during the standard MD simulations. The quantities are expressed as percentages over the total amount of frames. All simulations were performed using the HMR protocol and a time-step Δt=4 fs, except for the starred one, in which standard atomic masses and a time-step Δt=2 fs were used.

| $\text{Na}_v$ (SF) | System | Ions | Time (ns) | 0 (%) | 1 (%) | 2 (%) | 3 (%) | 4 (%) |
| --- | --- | --- | --- | --- | --- | --- | --- | --- |
| 1.2 (DEK<sup>+</sup>A)* | Model | [NaCl] | 500 | 7.7 | 66.3 | 25.1 | 0.9 | - |
| 1.1 (DEK<sup>+</sup>A) | Model | [NaCl] | 500 | 15.6 | 65.1 | 19.0 | 0.3 | - |
| 1.2 (DEK<sup>+</sup>A) | Model | [NaCl] | 500 | 10.9 | 59.7 | 28.3 | 1.1 | - |
| 1.6 (DEK<sup>+</sup>A) | Model | [NaCl] | 500 | 7.6 | 65.2 | 26.7 | 0.5 | - |
| 1.2 (DEK<sup>+</sup>A) | Cryo-EM | [NaCl] | 500 | 4.3 | 53.4 | 41.1 | 1.2 | - |
| 1.2 (DEK<sup>+</sup>A) | Cryo-EM | [NaCl] | 500 | 18.5 | 73.1 | 8.0 | 0.4 | - |

**Table 3:**
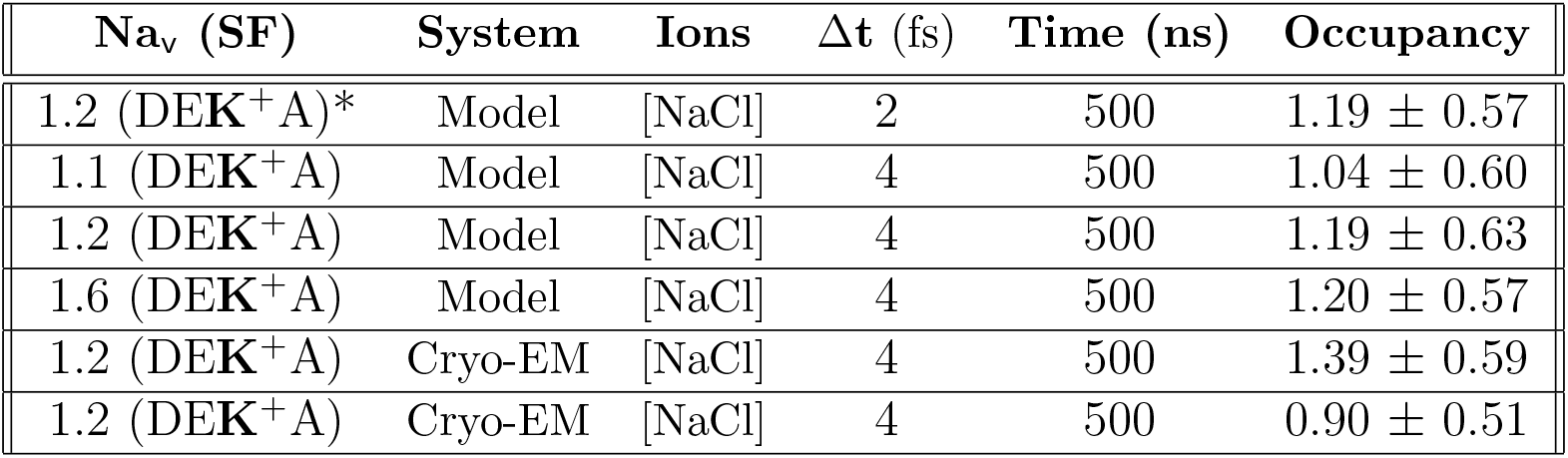
Average Na^+^ occupancy in the C/SF of the channels during standard MD simulations. The occupancy is introduced as average ± the standard deviation. All simulations used the HMR protocol and Δt=4 fs except for the starred one, where Δt=2 fs and employed standard masses.

We then looked at the specific positions of the Na^+^ ion in the C/SF in all the frames that showed a single cation permeating the filter. These snapshots form the majority of the ionic configurations revealed by our MD runs. To this aim, we combined the single-ion configurations from all six MD runs of the different systems. The aggregation of results from the various Na_v_s is justified by the fact that the structures are nearly identical in the filter region, a highly conserved domain of the Na_v_ family. In **Figure 5** we show the histograms of the *z* coordinate of the single Na^+^ ion in the C/SF (panel **A**), and of the nitrogen of K1422 (panel **B**), which is considered to play a relevant role in ionic permeation through mammalian Na_v_s.^46,86–91^

**Figure 5:**
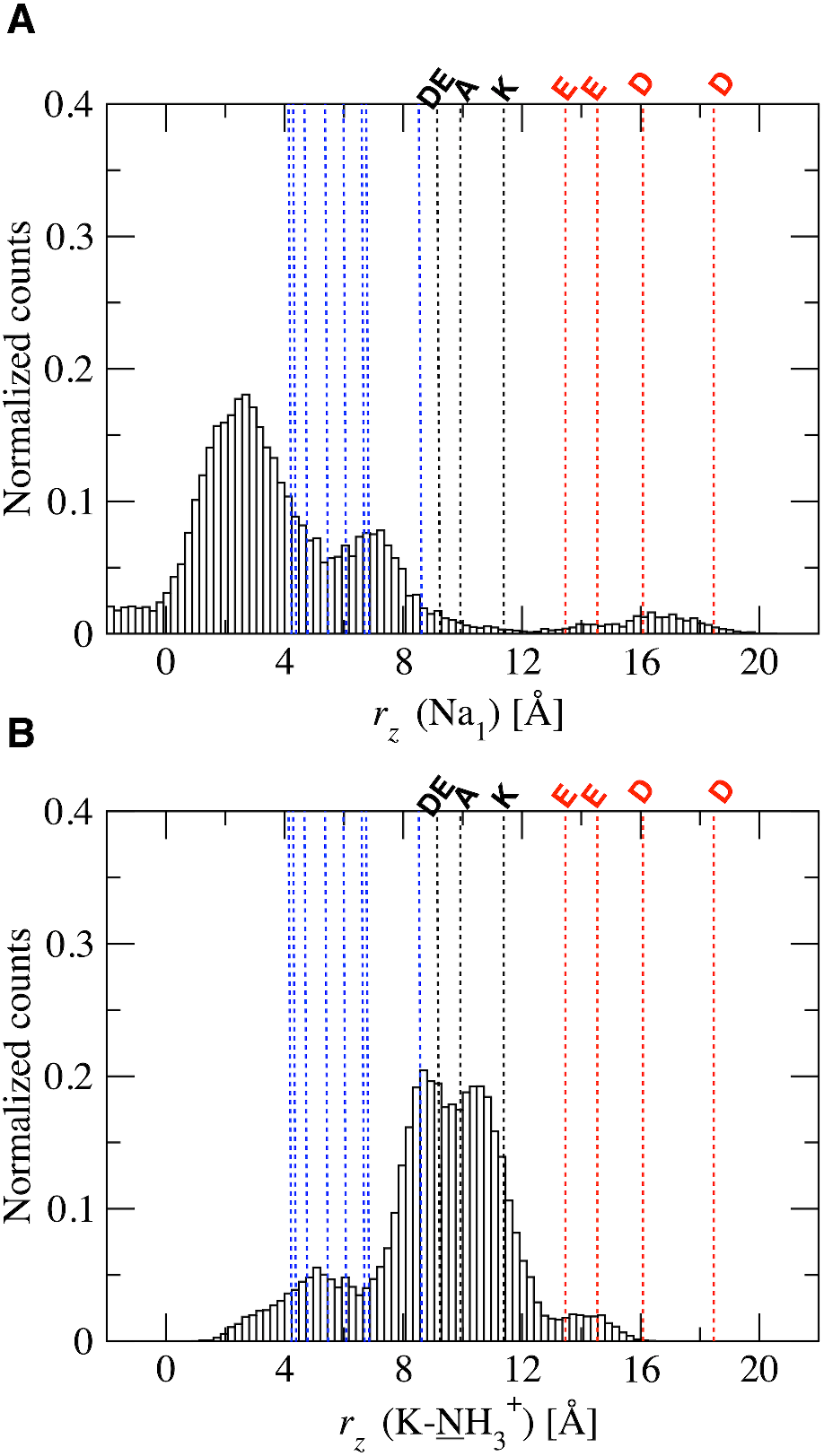
Histograms of the *z*-coordinate’s values of (**A**) a single Na^+^ ion in the C/SF and (**B**) the ammonium nitrogen of the DEKA ring. Vertical lines indicate the *z* coordinates of the C*α* atoms of C/SF residues from the equilibrated configuration of Na_v_1.2 (red for EEDD, black for DEKA and blue for each couple of residues right below the DEKA ring).

Results show that the single Na^+^ mostly occupies the lower portion of the SF, below the DEKA ring and in proximity of the inner rings of carbonyls. Conversely, the lysine side-chain resides at a higher position along the axis, where the positive charge is expected to interact with the negatively charged residues of the ring. These interactions involving the lysine might even occlude the SF, competing with the passage of other alkali ions from the extracellular environment. A set of six movies, showing different events of Na^+^ permeation throughout the C/SF and its residency in proximity of the lower rings, is provided as Supplementary Material.

Our findings of Na^+^ in proximity of the inner region of the SF are consistent with what observed in Ref. 93 via MD simulations of multiple Na_v_ channels (including 1.2, 1.4 and 1.7) starting from Cryo-EM structures.

Notably however, this position is different from the one occupied by a single Na^+^ ion in the Na_v_1.2 structure (PDB 6J8E). Yet, the Cryo-EM structure captures the channel with the C/SF occluded by the bound *μ*-conotoxin KIIIA, which limits a comparison with our computational investigations.

### 2D-FES calculations for sodium translocation through the Na_v_1.2 selectivity filter

We then used TAMD/OTFP simulations to reconstruct the thermodynamics of Na^+^ permeation through the C/SF of the Na_v_1.2 channel structures with different residue configurations, namely DEK^+^A, DEK^0^A, DEE^−^A, DEE^0^A (see Materials and Methods). We performed two sets of TAMD/OTFP simulations, named OTFP_1_ and OTFP_2_, that differ in the definition of the CVs and describe, respectively, single- and two-Na^+^ translocation.

In the next two paragraphs we show the average FESs obtained from multiple independent OTFP_1_ and OTFP_2_ runs, respectively, while results for the individual trajectories are reported in **Figures S5** to **S8**. All TAMD/OTFP runs are summarized in **Table S2**. In all simulations, the ions not involved in the CVs definition were excluded from the C/SF (see Material and Methods). In order to investigate the effect of this setting, we performed a set of independent simulations for the DEK^+^A system using a larger exclusion domain. Results are reported in **Figure S9** and do not show any remarkable difference with the smaller domain.

#### Single sodium ion translocation

For each of the DEK^+^A, DEK^0^A, DEE^−^A, DEE^0^A Na_v_1.2 systems, we ran six independent OTFP_1_ calculations, three starting from the homology model (**Figure S5**), and three starting from the Cryo-EM structure (**Figure S6**). Then, we calculated an average map by taking, for each point in CV space where the FESs are defined, the average of their FE values. The resulting averaged maps are reported in **Figure 6**, along with representative snapshots for the metastable and other states. Interestingly, the DEK^+^A system shows the broader low energy region, where it is possible to identify a gradually decreasing energy path for the ion through the SF and different configurations for the sidechain of K1422. Conversely, all other systems display a single isolated minimum associated with a unique orientation of the sidechain of residue 1422.

**Figure 6:**
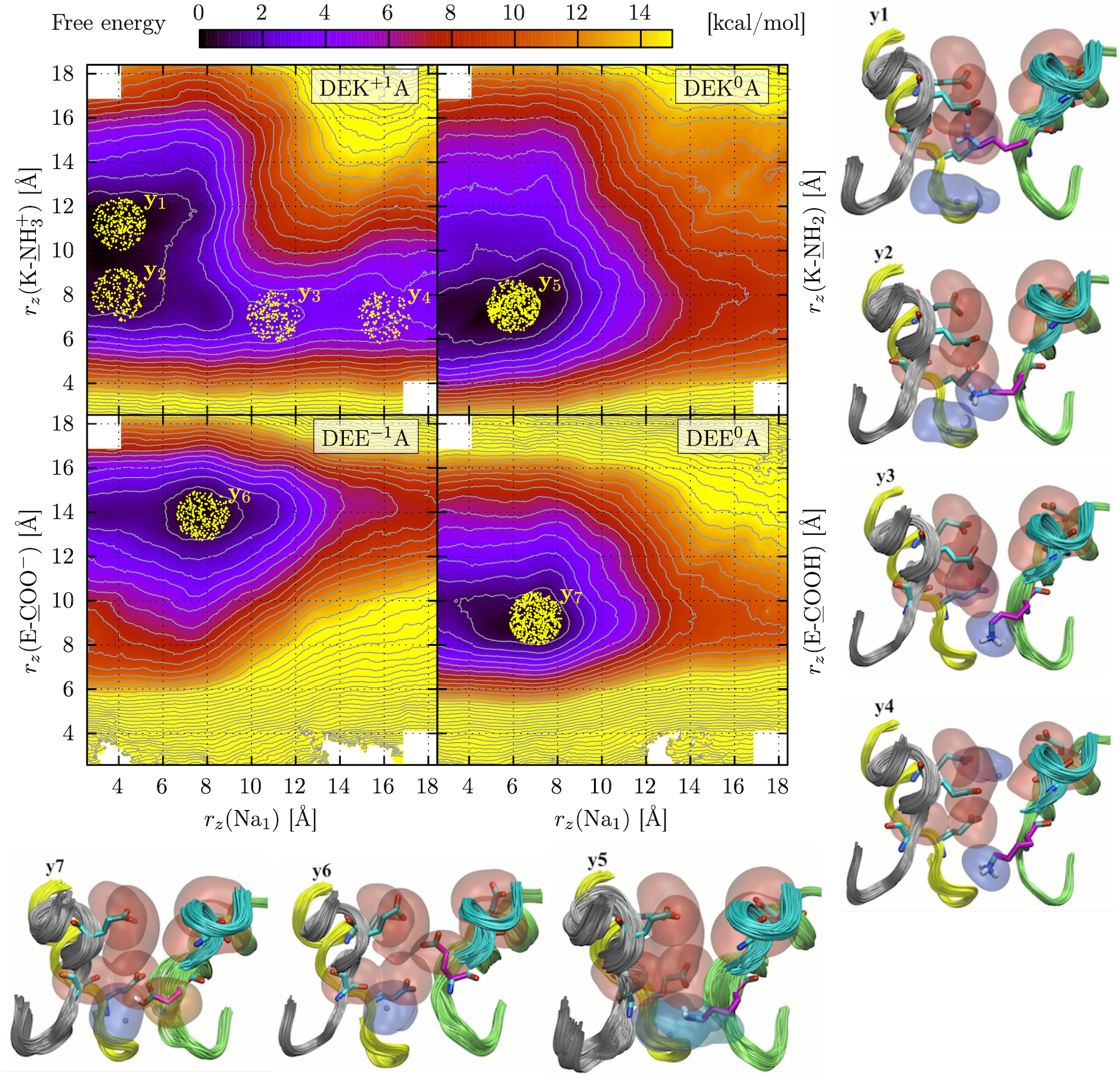
FESs for single Na^+^ translocation for each of the four systems, as indicated in the labels (DEK^+^A, DEK^0^A, DEE^−^A, DEE^0^A). Yellow dots indicate the sets of snapshots used for the representative configurations y_1_ to y_7_. For each set y_*i*_, the backbone of the various repeats is shown in ribbons, using the color code illustrated in **Figure 1**. A fragment of Domain IV (cyan) is not drawn for clarity. The transparent surfaces represent the volume occupancy of Na^+^ and the tip atoms of the DEKA and EEDD sidechains, and they are colored according to atomic charge (red for negative and blue for positive charges). The surface of residue 1422 is colored cyan for DEK^0^A and orange for DEE^0^A).

For the DEK^+^A system, the positions of the single Na^+^ and of K1422 at the energy minima are well correlated with the high probability locations obtained from our standard MD simulations (see **Figure 5**), and with what described in Ref. 93. In these configurations (see the states y_1_ and y_2_ in **Figure 6**), the Na^+^ is located at the inner portion of the SF (*r_z_*(Na_1_) ~ 4 Å), in the proximity of the internal carbonyl oxygen atoms belonging to the two residues underneath the DEKA ring. The K1422 ammonium group is found either at the level of the DEKA backbone (at 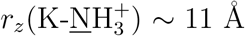, y_1_) or at a lower position (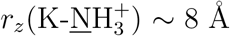, y_2_). Using the string method,^137–139^ we computed the minimum free energy path (MFEP) over the DEK^+1^A surface, shown as a white trace over the corresponding map in **Figure 7**, where we also report the FE profile along the path. The MFEP allows us to reconstruct the following mechanism for single ion permeation: Na^+^ access to the upper part of the C/SF is possible only when K1422 is in a lower position, interacting with the upper ring of the internal carbonyls (configurations y_4_ and y_3_); once the ion reaches its energy minimum below the DEKA ring, K1422 swings in an upper position, at the level of the DEKA backbone (state y_1_). This picture is consistent with the role of the protonated K1422 suggested in Ref.s 70,140, where it is proposed that the charged lysine should interact with the negatively charged side chain of the DEKA ring to block the access of ions from the extracellular space.

**Figure 7:**
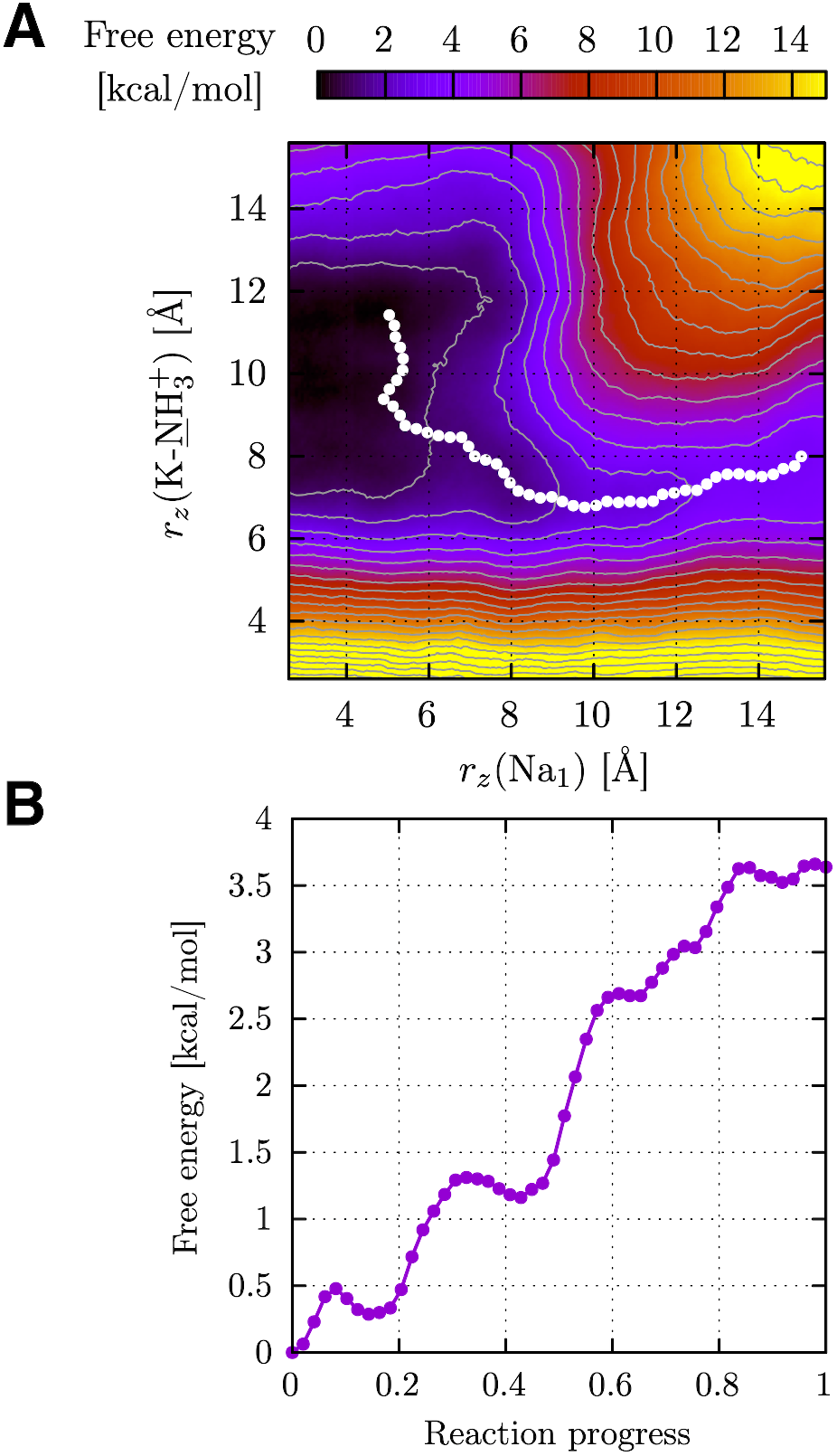
**(A)** MFEP (white dots) of the DEK^+1^A system’s averaged FES, superimposed on the corresponding map. (**B**) FE profile along the MFEP.

Opposite to the DEK^+1^A configuration, all the other systems’s FESs show an isolated minimum, again with the ion located in the region surrounded by the carbonyl oxygen atoms below the DE*A ring (where * indicates the variable residue among the different configurations), at 4 Å < *r_z_*(Na_1_) < 8 Å. In all three cases, the modification of the K1422 residue results in higher energy values in the upper part of the C/SF. The system with the DEK^0^A ring is characterized by a single metastable state in the FES, y_5_ (**Figure 6)**, defined by the Na^+^ ion at *r_z_*(Na_1_) ~ 6 Å and the *r_z_*(K-<u>N</u>H_2_) at ~ 7 Å. Other works investigated the role of different protonation states of the DEKA lysine^46,70^ in ionic permeation. While extensive unbiased MD simulations of a chimeric bacterial/eukaryotic channel^46^ suggested that the uncharged lysine plays a minor role in the selectivity process, Monte Carlo simulations on multiple eukaryotic structures^70^ proposed that the uncharged lysine is responsible of escorting the Na^+^ ion to the inner carbonyl rings, resulting in a configuration that is very similar to the one identified in our DEK^0^A TAMD/OTFP simulations (state y_5_).

Also the K1422E mutation (either DEE^−^A or DEE^0^A) affects the thermodynamic features of Na^+^ translocation. In the DEE^−^A minimum (y_6_), the Na^+^ ion is localized at *r_z_*(Na_1_) ~ 8 Å, with the E1422 sidechain confined in the upper region of the SF, at *r_z_*(E-<u>C</u>OO^−^) ~ 14 Å. In the case of DEE^0^A, the FE minimum (y_7_) is located at *r_z_*(Na_1_) ~ 7 Å and *r_z_*(E-<u>C</u>OOH) ~ 9 Å i.e. with the E1422 CD atom in the proximity of the inner ring’s backbone, in a configuration similar to that of the DEK^0^A system (y_5_), but with the ion and the glutamate sidechain in a slightly higher position.

#### Two sodium ions translocation

We then computed 2D FESs using the OTFP_2_ set-up of CVs illustrated in Materials and Methods, i.e. by considering as CVs the vertical position of the COM of two ions and of the sidechain of residue 1422. Indeed, multiple Na^+^ ion configurations were observed in a chimeric bacterial/mammalian model including the Na_v_1.2 filter, ^46^ and extensively discussed in many works presenting experimental results. ^80–83^

As detailed in **Table S2**, and similarly to what described for the OTFP_1_ simulations, for each (DEK^+^A, DEK^0^A, DEE^−^A, DEE^0^A) system we performed six independent OTFP_2_ calculations, three using the homology model (**Figure S7**), and three the Cryo-EM structure (**Figure S8**), and averaged the results in a single map.

The final averaged maps are reported in **Figure 8**, along with a set of representative snapshots for the metastable and other states. The DEK^+^A FES shows a shallow (z_1_) and two deep FE minima (z_2_ and z_3_). The z_1_ configuration is characterized by the two ions at the inner position, with their COM_*z*_ at ~ 8 Å and the lysine in proximity of the EEDD ring, nearly at the upper limit of the sampled space, 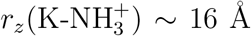. The z_2_ and z_3_ states correspond to higher positions for the COM_*z*_ of the ions and lower positions for K1422, i.e. COM_*z*_ (Na_1_, Na_2_) ~ 10 Å, and 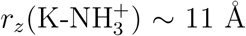, for z_2_, and COM_*z*_ (Na_1_, Na_2_) ~ 14 Å and 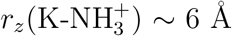, for z_3_. Hence, by considering the sequence of states in the order z_3_ to z_1_, we can describe the movement of 2 Na^+^ ions within the C/SF as follows:

- z_3_: the two ions are at their highest positions, while K1422 is at its lowest, and the sequence of positive charges from the extracellular side to the inner part of the filter is [Na^+^, Na^+^, 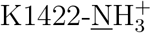].
- z_2_: the ions move to a lower position (in particular the inner one), and K1422 switches to an intermediate one, between them; the sequence of positive charges from the extracellular side to the inner region is now [Na^+^, 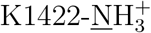, Na^+^].
- z_1_: from the previous state, a less stable configuration is accessible, characterized by a slightly lower position of the ions, and a considerably higher location of K1422; the sequence of positive charges is now [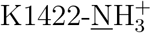, Na^+^, Na^+^].

**Figure 8:**
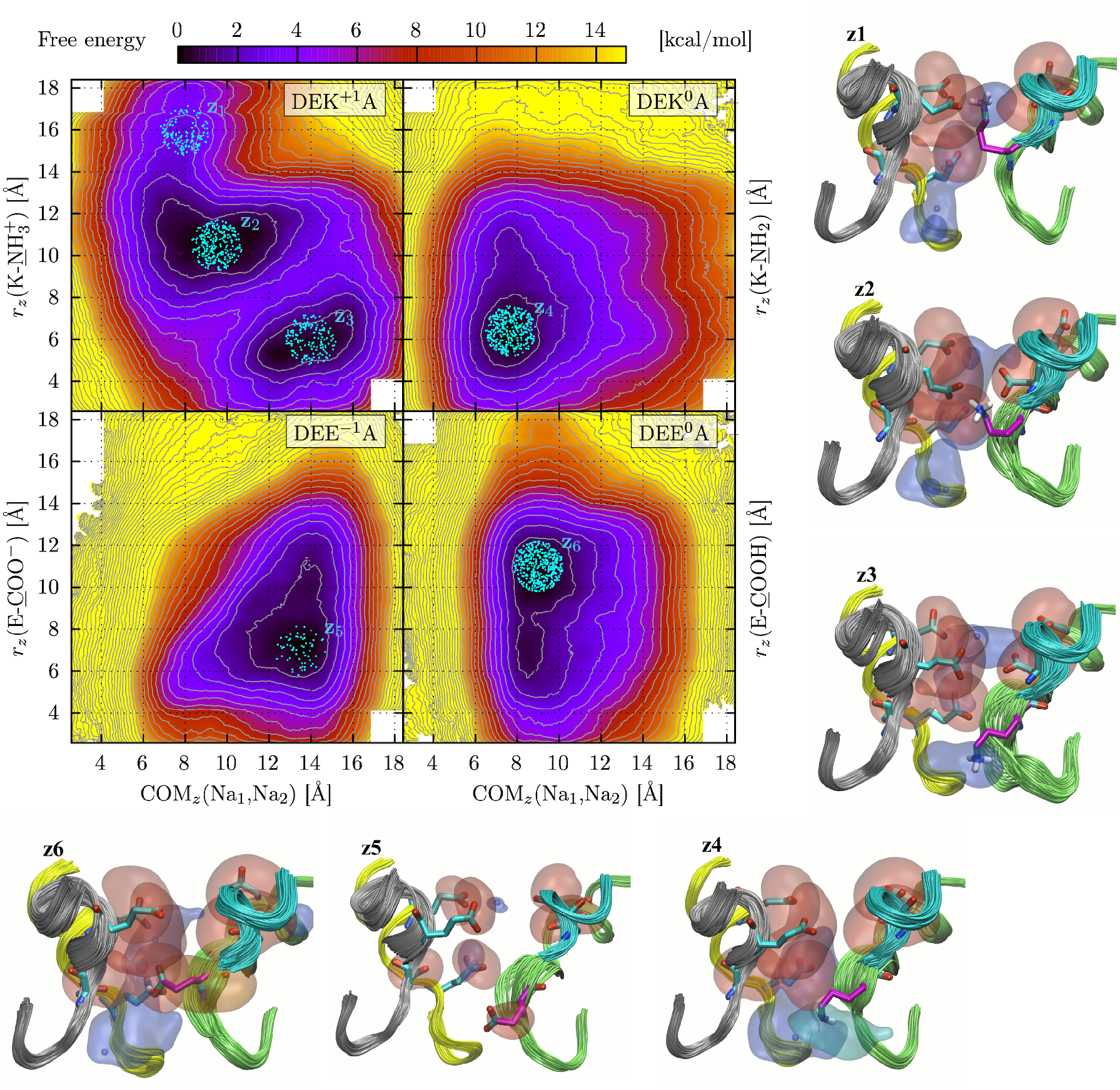
FESs for two Na^+^ ion translocation in the SF of each of the four systems, as indicated in the labels (DEK^+^A, DEK^0^A, DEE^−^A, DEE^0^A). Cyan dots indicate the snapshots used for the representative configurations z_1_ to z_6_, which are reported as explained in **Figure 6**.

These multiple features are completely lost in the maps of the other three systems DEK^0^A, DEE^−^A and DEE^0^A. The DEK^0^A FES shows a single minimum, z_4_, defined by COM_*z*_ (Na_1_, Na_2_) ~ 8 Å and *r_z_*(K-<u>N</u>H2) ~ 6 Å, i.e. with the ions and the uncharged lysine at lower positions. This configuration of the side-chain is again similar to that observed in Ref. 70. When the DEKA filter is replaced by DEE^−^A, the two ions are substantially trapped in the upper part of the SF (state z_5_, COM_*z*_(Na_1_, Na_2_) at ~ 14 Å with 1 ion at the level of the EEDD motif and 1 ion in proximity of the DEEA ring), and their access to the inner region is precluded. In the case of the DEE^0^A system, the main metastable configuration (z_6_) corresponds to COM_*z*_ (Na_1_, Na_2_) ~ 9 Å and *r_z_*(E-<u>C</u>OOH) ~ 11 Å, with a less populated state with the uncharged glutamate at a lower position. This latter state is similar to z_4_, revealing again similarities among the two uncharged systems (DEK^0^A and DEE^0^A).

## Conclusions

Voltage gated sodium channels play a pivotal role in the regulation of excitable cells, and mutations in genes encoding them have been linked to numerous disorders affecting nervous system function, heart rhythm and muscle contraction. Therefore, understanding the structural and mechanistic features of the cationic fluxes through the Na_v_ conductivity/selectivity filter (C/SF) is essential to accelerate the development of new efficient pharmacological treatments. In this study, we used a combination of molecular modeling, standard MD and enhanced sampling to describe in detail the structural features of ion permeation through the SF of human neuronal Na_v_ channels. We combined results from MD simulations of homology-based models of Na_v_1.1, 1.2 and 1.6 and the recent Cryo-EM Na_v_1.2 structure (PDB ID: 6J8E) to describe the thermodynamics of Na^+^ residency within the C/SF. We ran simulations on multiple variants of the key SF DEKA ring, by modifying the positively charged K (K1422 in Na_v_1.2), which is demonstrated to be essential for selectivity. Our simulations include a set of enhanced sampling MD simulations employing the TAMD/OTFP method to reconstruct 2D FESs for single and double Na^+^ ions through the C/SF of the Na_v_1.2 structures. TAMD/OTFP has been previously assessed as an efficient tool to investigate systems with hidden FE barrier. FESs were obtained separately for two CV sets, the vertical position of one or two Na^+^ ions in the C/SF and of the sidechain of residue 1422. We also investigated the role of K1422 protonation and of charged and uncharged K1422E mutants. The obtained FESs show that in the WT, DEK^+^A system there is a pathway through the C/SF with low FE values that the ions can follow to reach the inner portion of the filter, both as single or two ions configurations. For all other systems (DEKA, DEEA, DEE^−^A), the FESs display single, isolated minima, mostly localized in the inner portion of the SF. Our results confirm the active role of the lysine residue in controlling Na^+^ transport, describing possible 1 and 2-ion occupancy states, as those investigated in the long standard MD simulations described in Ref. 46, where the authors used a modified structure of a bacterial Na_v_ channel by including the human DEKA motif. To the best of our knowledge, our simulations provide the first reconstruction of the thermodynamics of multi Na^+^ permeation through Na_v_ 3D configurations based on the recent Cryo-EM resolved structures. The present approach, leveraging the TAMD/OTFP methodology, may be used in further works to extend the space of sampling and, in addition, to explore the selectivity mechanisms of neuronal Na_v_ channels.

## Main abbreviations and notations

CD: cross distance
C/SF: conductivity/selectivity filter
Cryo-EM: cryogenic electron microscopy
DEKA: notation for the Asp, Glu, Lys, Ala residues in the inner DEKA ring of the selectivity filter
EEDD: notation for the Asp, Asp, Glu, Glu residues in the outer EEDD ring
FF: force field
FE: free energy
MD: molecular dynamics
Na_v_: voltage gated sodium channel
OTFP: on-the-fly free-energy parametrization
PD: pore domain
POPC: 1-palmitoyl-2-oleoyl-sn-glycero-3-phosphocholine
RMSD: root mean standard deviation
S1-S6: labels for the six *α* helices in each domain of a human voltage gated sodium Na_v_ channel
TAMD: temperature accelerated molecular dynamics
SF: selectivity filter

## Associated Contents

The Supporting Information is available

- Representation of the Na_v_ 1.1 and Na_v_ 1.6 SFs. Superposition between the Cryo-EM resolved Na_v_1.2 structure (PDB ID: 6J8E) and the homology based Na_v_1.2 model. Backbone RMSD calculations for the standard MD simulations. Cross distance analysis. Individual FESs for OTFP_1_ and OTFP_2_ (homology-based Na_v_1.2 structure). Individual FESs for OTFP_1_ and OTFP_2_ (Cryo-EM resolved Na_v_1.2 structure - PDB ID: 6J8E). Average OTFP_1_ and OTFP_2_ for the homology-based Na_v_1.2 structure in the presence of the extended exclusion rectangle. **PDF**
- List of movies. **MP4**

## Author Contributions

- **Conceptualization:** Luca Maragliano and Cameron F. Abrams.
- **Data curation:** Giulio Alberini, S. Alexis Paz, and Beatrice Corradi.
- **Formal analysis:** S. Alexis Paz, Giulio Alberini, and Beatrice Corradi.
- **Funding acquisition and resources:** Luca Maragliano, Fabio Benfenati and S. Alexis Paz.
- **Investigation:** Giulio Alberini, S. Alexis Paz, Beatrice Corradi, Luca Maragliano and Cameron F. Abrams.
- **Project administration:** Giulio Alberini, Luca Maragliano.
- **Supervision:** Luca Maragliano and Cameron F. Abrams.
- **Visualization:** S. Alexis Paz, Giulio Alberini and Beatrice Corradi.
- **Writing - original draft:** Giulio Alberini.
- **Writing - review & editing:** S. Alexis Paz, Giulio Alberini, Beatrice Corradi, Luca Maragliano, Fabio Benfenati and Cameron F. Abrams.

## Acknowledgments

We thank Alessia Vignolo, Mattia Pini and Sergio Decherchi for the kind assistance at the Italian Institute of Technology (IIT) computing center. We are grateful to Alessandro Berselli, Diego Moruzzo and Andrea L. Benfenati for useful help and technical assistance. We acknowldge Matteo Ceccarelli, Pasquale Striano and Federico Zara for very useful discussions.

Computing time allocations were granted by the CINECA supercomputing center under the ISCRA initiative. We also gratefully acknowledge the HPC infrastructure and the Support Team at Fondazione Istituto Italiano di Tecnologia.

This work also used computational resources from CCAD – Universidad Nacional de Córdoba (https://ccad.unc.edu.ar/), which are part of SNCAD – MinCyT, República Argentina. Special thanks to CCAD for giving us priority and exclusive access to the computer resources in the begining of this project. The study was supported by research grants from the Compagnia di San Paolo Torino (2015.0546 and 2019.34760 to FB); IRCCS Ospedale Policlinico San Martino (Ricerca Corrente and 5 × 1000 to LM and FB) and Italian Ministry of University and Research (2017-A9MK4R and 2020-WMSNBL to FB). The support of Telethon-Italy (Grant GGP19120 to FB) is also acknowledged. S. A. Paz acknowledges the financial support from the Agencia Nacional de Promoción Científica y Tecnológica (ANPCyT-FONCyT Grant PICT-2017-0621).

## Ethics Statements

In this work, no animal or human study is presented.

