## Supplementary information for "Molecular Dynamics Simulations of Ion Permeation in human NaV Channels"

¶*Universidad Nacional de Córdoba. Facultad de Ciencias Químicas. Departamento de Química Teórica y Computacional. Córdoba (X5000HUA), Argentina.*

§*Consejo Nacional de Investigaciones Científicas y Técnicas (CONICET), Instituto de Fisicoquímica de Córdoba (INFIQC), Córdoba (X5000HUA), Argentina.*

∥*Department of Experimental Medicine, Università degli Studi di Genova, Viale Benedetto XV, 3, 16132, Genova, Italy.*

⊥*Department of Chemical and Biological Engineering, Drexel University, Philadelphia, PA 19104, United States.*

#*Department of Life and Environmental Sciences, Polytechnic University of Marche, Via Brecce Bianche, 60131, Ancona, Italy.*

[Δ] Giulio Alberini and S. Alexis Paz should be regarded as joint first authors.

#### Definition of the Cross Distances

In this section, we report the list of the cross distances (CDs) mapped during the standard MD simulations. The average for each CD is reported in **Table S3**.

1. EEDD motif

   - **d1A**: E387CA - D1426CA; **d1B**: E387CD - D1426CG
   - **d2A**: E945CA - D1717CA; **d2B**: E945CD - D1717CG

| Residue numbering in the EEDD domain (d1-d2) | | | | |
|---|---|---|---|---|
| **System** | **Rep. I** | **Rep. II** | **Rep. III** | **Rep. IV** |
| NaV1.4 | 409 | 764 | 1248 | 1539 |
| NaV1.1 | 385 | 954 | 1436 | 1727 |
| **NaV1.2** | **387** | **945** | **1426** | **1717** |
| NaV1.6 | 373 | 939 | 1417 | 1708 |

2. DEKA motif

   - **d3A** D384CA - K1422CA; **d3B** D384CG - K1422CE
   - **d4A** E942CA - A1714CA; **d4B** E942CD - A1714CB

| Residue numbering in the DEKA domain (d3-d4) | | | | |
|---|---|---|---|---|
| **System** | **Rep. I** | **Rep. II** | **Rep. III** | **Rep. IV** |
| NaV1.4 | 406 | 761 | 1244 | 1536 |
| NaV1.1 | 382 | 951 | 1432 | 1724 |
| **NaV1.2** | **384** | **942** | **1422** | **1714** |
| NaV1.6 | 370 | 936 | 1413 | 1705 |

3. BACKBONE scaffold

- **d5A** COM between T382CA and Q383CA - COM between T1420CA and F1421CA; **d5B** COM between T382O and Q38O - COM between T1420O and F1421O
- **d6A** COM between C940CA and G941CA - COM T1712CA and S1713CA; **d6B** COM between C940O and G941O - COM T1712O and S1713O

| Residue numbering (d5-d6) | | | | |
|---|---|---|---|---|
| **System** | **Rep. I** | **Rep. II** | **Rep. III** | **Rep. IV** |
| NaV1.4 | T404-Q405 | C759-G760 | T1242-F1243 | T1534-S1535 |
| NaV1.1 | T380-Q381 | C949-G950 | T1430-F1431 | T1722-S1723 |
| **NaV1.2** | **T382-Q383** | **C940-G941** | **T1420-F1421** | **T1712-S1713** |
| NaV1.6 | T368-Q369 | C934-G935 | T1411-F1412 | T1703-S1704 |

- **d7A** L421CA-I1469CA; **d7B** L421CG-I1469CB
- **d8A** F978CA - I1775CA; **d8B** COM among the carbon atoms of the F978 aromatic ring - I1775CB
- **d9A** F978CA - Y1771CA; **d9B** COM among the carbon atoms of the F978 aromatic ring - COM among the carbon atoms of the Y1771 aromatic ring

| Residue numbering (d7-d8-d9) | | | | |
|---|---|---|---|---|
| **System** | **Rep. I** | **Rep. II** | **Rep. III** | **Rep. IV** |
| NaV1.4 | L443 | F797 | I1291 | I1597-Y1593 |
| NaV1.1 | L419 | F987 | I1479 | I1785-Y1781 |
| **NaV1.2** | **L421** | **F978** | **I1469** | **I1775-Y1771** |
| NaV1.6 | L407 | F972 | I1460 | I1765-Y1761 |

### Supplementary Numbered Tables

**Table S1:** Summary of the standard MD simulations with a 150 mM NaCl concentration. In all the runs the complete configuration (Conf.) included both the VSDs and the central pore.

| **$Na_v$ (C/SF)** | **Conf.** | **System** | **Method** | **$\Delta$t** (fs) | **Time (ns)** |
|---|---|---|---|---|---|
| 1.2 (DE**K**$^+$A) | VSDs+Pore | Model | MD | 2 | 500 × 1 |
| 1.1 (DE**K**$^+$A) | VSDs+Pore | Model | MD | 4 | 500 × 1 |
| 1.2 (DE**K**$^+$A) | VSDs+Pore | Model | MD | 4 | 500 × 1 |
| 1.6 (DE**K**$^+$A) | VSDs+Pore | Model | MD | 4 | 500 × 1 |
| 1.2 (DE**K**$^+$A) | VSDs+Pore | Cryo-EM | MD | 4 | 500 × 2 |

**Table S2:** Summary of the TAMD-OTFP simulations with a 150 mM NaCl concentration. (*) indicates simulations that were performed in bigger exclusion rectangle that have vertices (-6, -6, -4) and (6, 6, 22) Å instead of (-6, -6, -4) and (6, 6, 18) Å . All simulations were performed with a time-step $\Delta$t = 2 fs.

| **Na$_v$ (SF)** | **K/E1422** | **Conf.** | **System** | **Method** | **Time (ns)** | **-** |
|---|---|---|---|---|---|---|
| 1.2 (DE**K**$^+$A) | **charged** | Pore | Model | OTFP$_1$ | 100 × 3 | |
| 1.2 (DE**K**$^+$A) | **charged** | Pore | Model | OTFP$_2$ | 100 × 3 | |
| 1.2 (DE**K**$^+$A) | **charged** | Pore | Model | OTFP$_1$ | 100 × 3 | * |
| 1.2 (DE**K**$^+$A) | **charged** | Pore | Model | OTFP$_2$ | 100 × 3 | * |
| 1.2 (DE**K**$^0$A) | **uncharged** | Pore | Model | OTFP$_1$ | 100 × 3 | |
| 1.2 (DE**K**$^0$A) | **uncharged** | Pore | Model | OTFP$_2$ | 100 × 3 | |
| 1.2 (DE**E**$^-$A) | **charged** | Pore | Model | OTFP$_1$ | 100 × 3 | |
| 1.2 (DE**E**$^-$A) | **charged** | Pore | Model | OTFP$_2$ | 100 × 3 | |
| 1.2 (DE**E**$^0$A) | **uncharged** | Pore | Model | OTFP$_1$ | 100 × 3 | |
| 1.2 (DE**E**$^0$A) | **uncharged** | Pore | Model | OTFP$_2$ | 100 × 3 | |
| 1.2 (DE**K**$^+$A) | **charged** | Pore | Cryo-EM | OTFP$_1$ | 125 × 3 | |
| 1.2 (DE**K**$^+$A) | **charged** | Pore | Cryo-EM | OTFP$_2$ | 125 × 3 | |
| 1.2 (DE**K**$^0$A) | **uncharged** | Pore | Cryo-EM | OTFP$_1$ | 125 × 3 | |
| 1.2 (DE**K**$^0$A) | **uncharged** | Pore | Cryo-EM | OTFP$_2$ | 125 × 3 | |
| 1.2 (DE**E**$^-$A) | **charged** | Pore | Cryo-EM | OTFP$_1$ | 125 × 3 | |
| 1.2 (DE**E**$^-$A) | **charged** | Pore | Cryo-EM | OTFP$_2$ | 125 × 3 | |
| 1.2 (DE**E**$^0$A) | **uncharged** | Pore | Cryo-EM | OTFP$_1$ | 125 × 3 | |
| 1.2 (DE**E**$^0$A) | **uncharged** | Pore | Cryo-EM | OTFP$_2$ | 125 × 3 | |

**Table S3:** Summary of the cross distances measured in the Cryo-EM $Na_v1.2$ PDB: 6J8E (second column) and as the average among the mean values of the single standard MD simulations (third column).

| Cross Distance | PDB ID:6J8E | $\mu \pm \sigma$ |
|---|---|---|
| d1A | 17.9 | $17.06 \pm 0.37$ |
| d1B | 13.2 | $12.50 \pm 0.40$ |
| d2A | 17.6 | $19.37 \pm 1.04$ |
| d2B | 12.0 | $13.88 \pm 1.01$ |
| d3A | 10.8 | $10.45 \pm 0.64$ |
| d3B | 9.1 | $8.60 \pm 0.57$ |
| d4A | 11.9 | $12.13 \pm 0.90$ |
| d4B | 10.2 | $10.94 \pm 0.81$ |
| d5A | 11.3 | $12.93 \pm 0.43$ |
| d5B | 11.2 | $9.41 \pm 0.35$ |
| d6A | 11.6 | $14.21 \pm 1.29$ |
| d6B | 12.9 | $10.21 \pm 0.62$ |
| d7A | 13.7 | $13.30 \pm 1.07$ |
| d7B | 11.1 | $9.79 \pm 1.01$ |
| d8A | 15.7 | $15.94 \pm 1.69$ |
| d8B | 11.6 | $14.01 \pm 2.03$ |
| d9A | 15.7 | $14.11 \pm 0.50$ |
| d9B | 12.3 | $9.14 \pm 0.43$ |

### Supplementary Figures

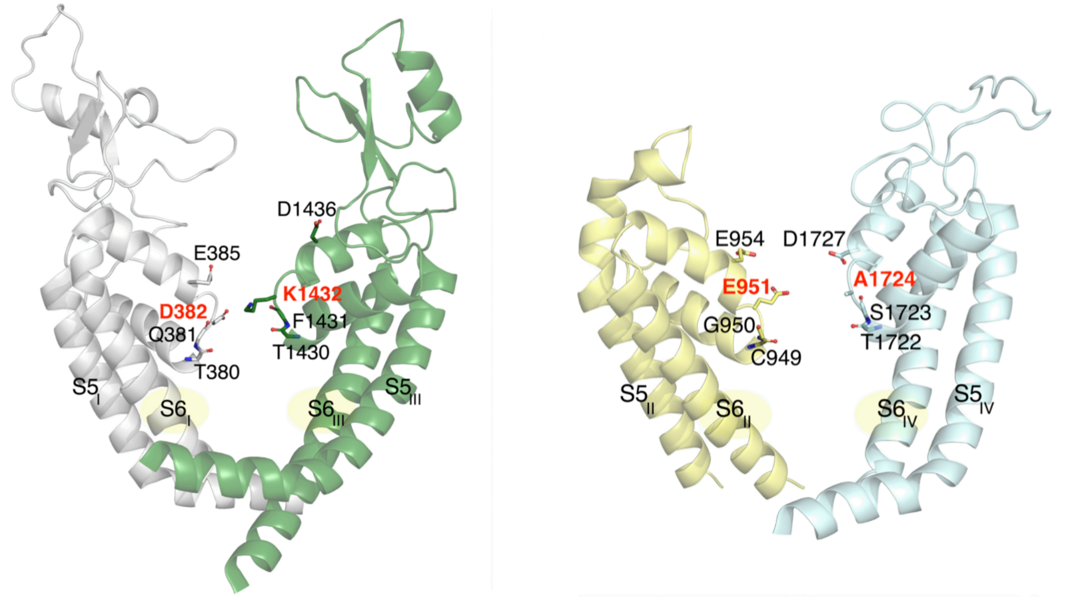


**Figure S1:** Representation of the asymmetric $Na_v1.1$ C/SF. The residues of the external $E_IE_{II}D_{III}D_{IV}$ motif, above the DEKA domain, are also shown.

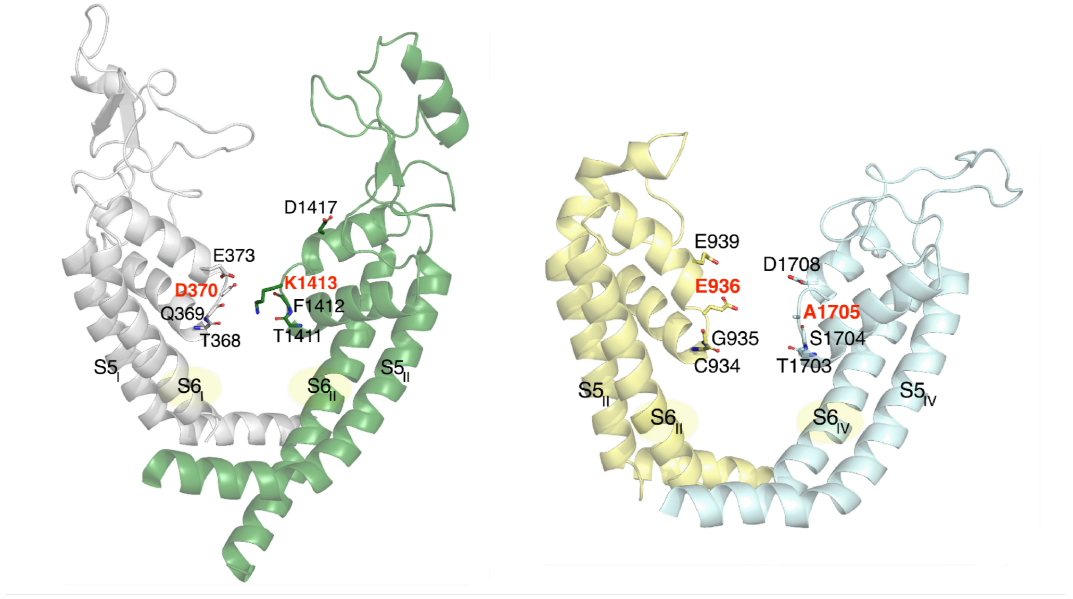


**Figure S2:** Representation of the asymmetric $Na_v1.6$ C/SF. The residues of the external $E_IE_{II}D_{III}D_{IV}$ motif, above the DEKA domain, are also shown.

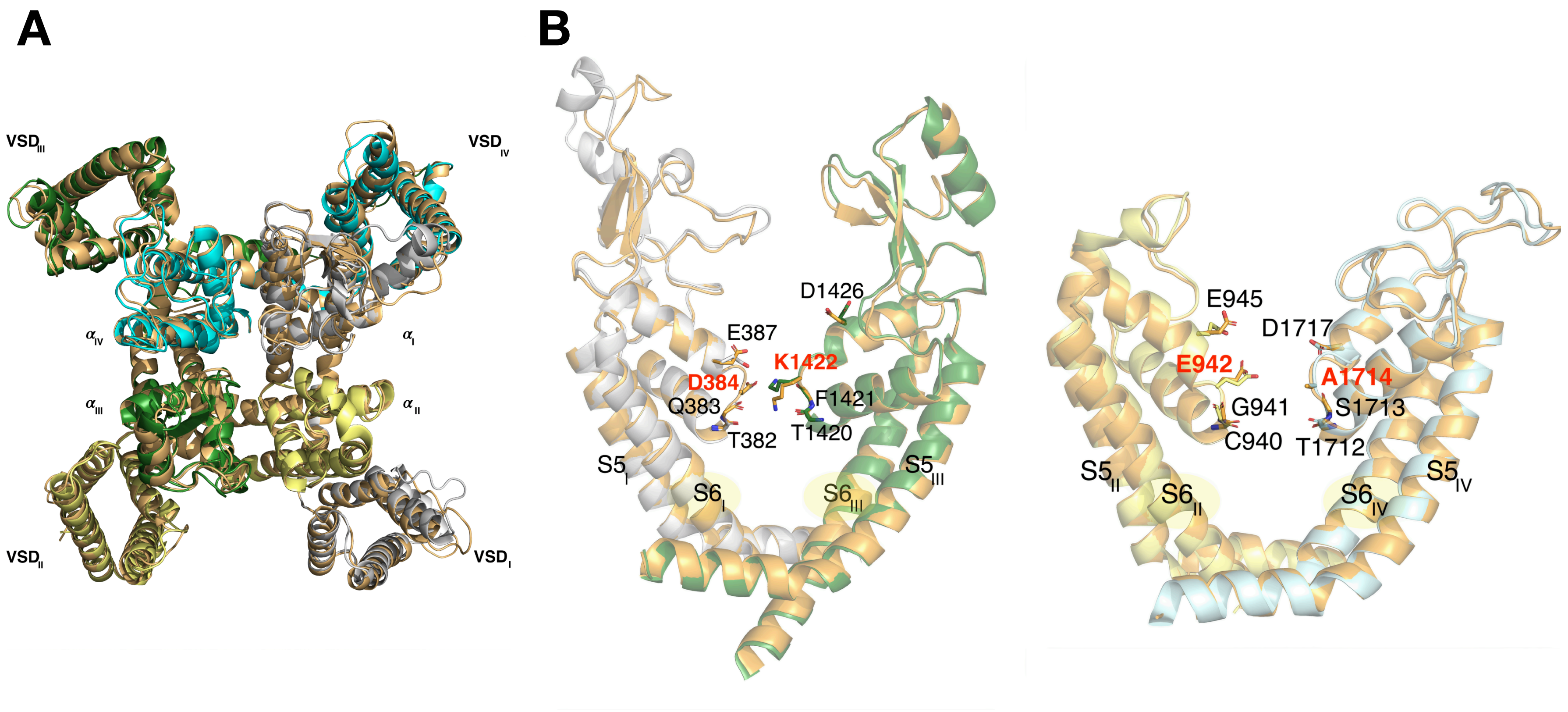


**Figure S3:** Superposition between the Cryo-EM resolved Na$_v$1.2 structure (PDB ID: 6J8E), in brown, and the homology-based model of the same channel, represented with the same legend of Figure 1: domain I, gray; domain II, yellow; domain III, green; domain IV, cyan. **A.** Extracellular view of the two superimposed structure. **B.** Representation of the asymmetric Na$_v$1.2 C/SF in the two configurations.

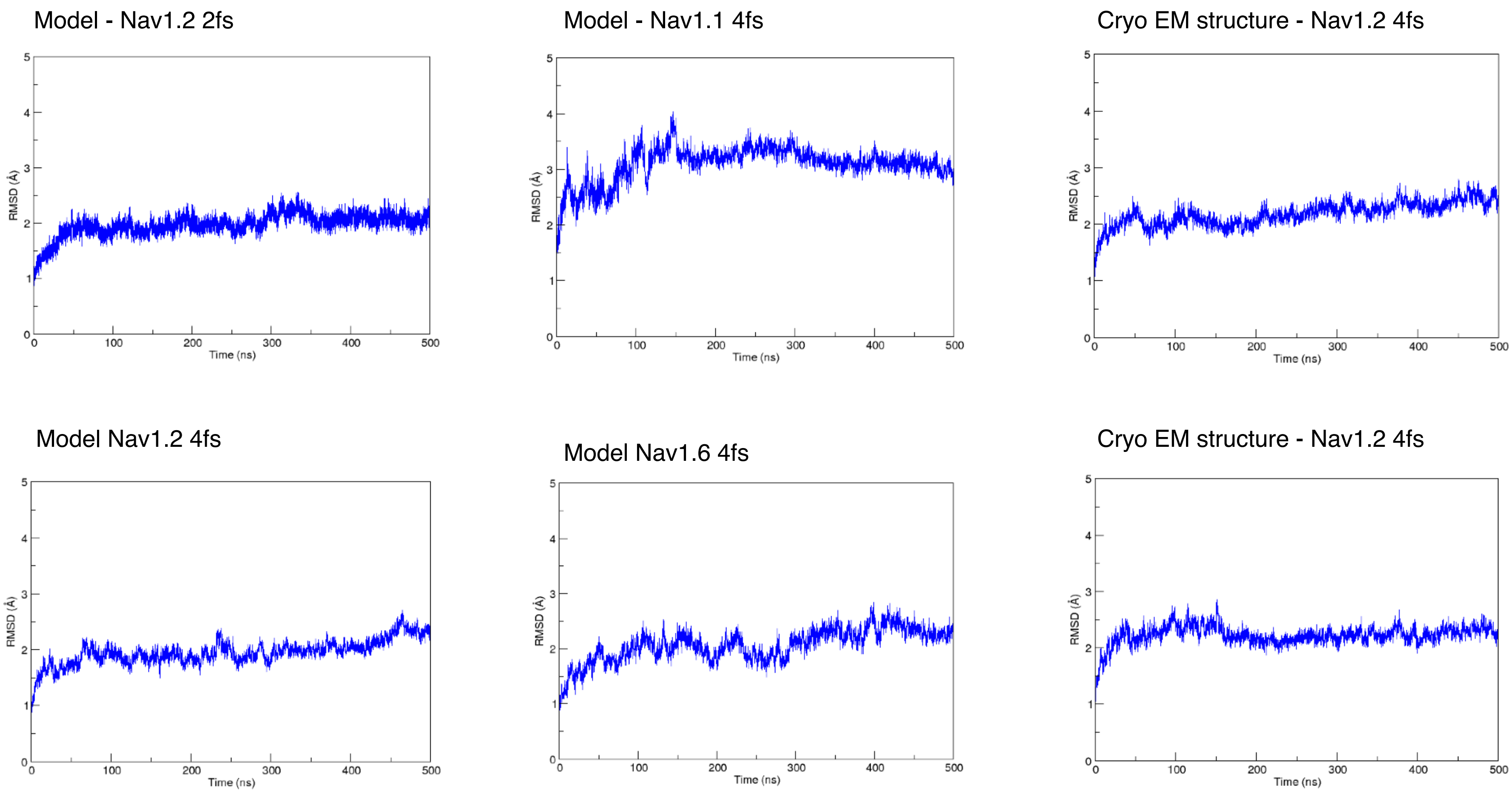


**Figure S4:** Backbone RMSD of the pore region for each system during standard MD simulations.

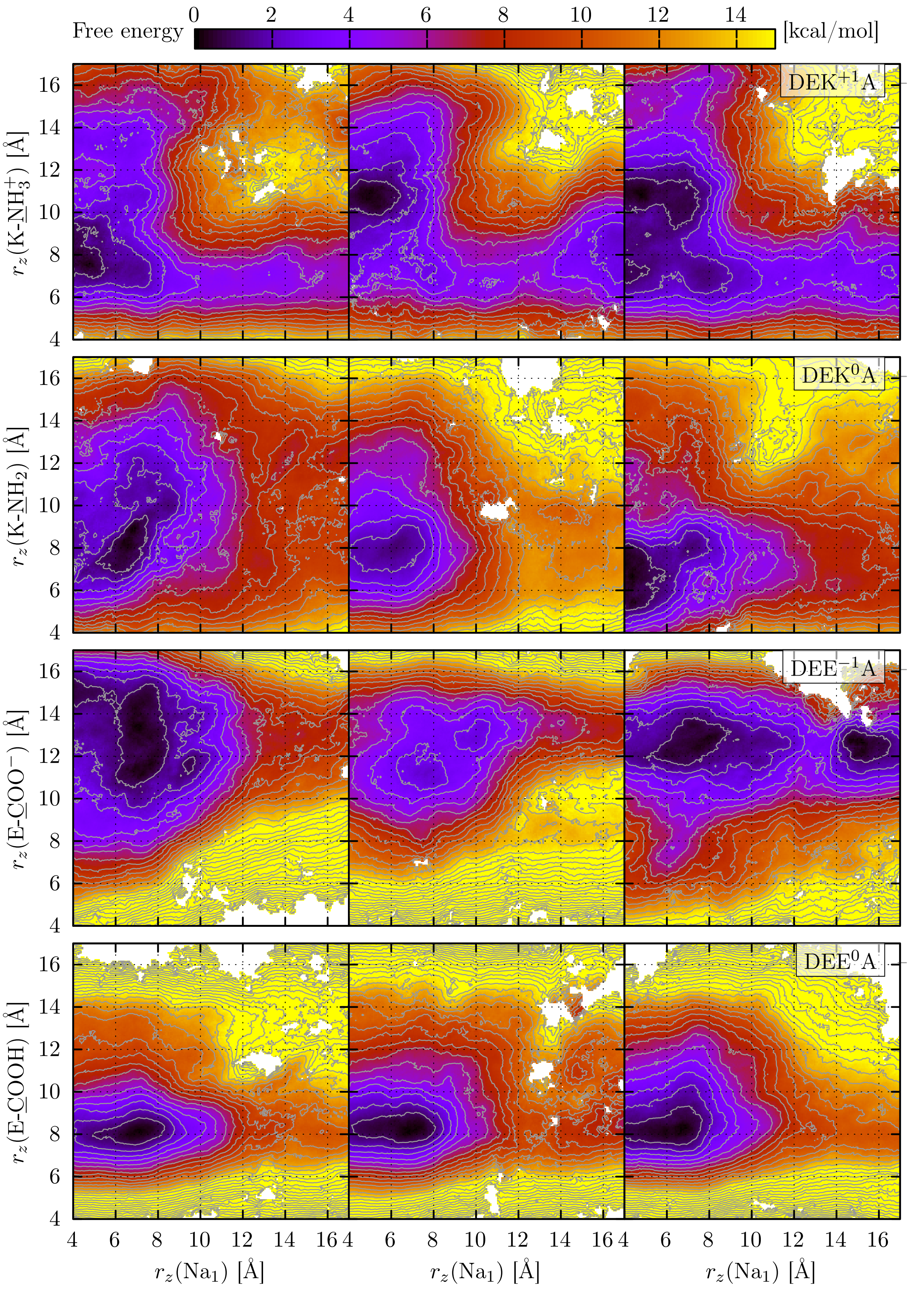


**Figure S5:** OTFP$_1$ FESs for single $Na^+$ translocation for each of the four systems of the Na$_v$1.2 model, as indicated in the labels.

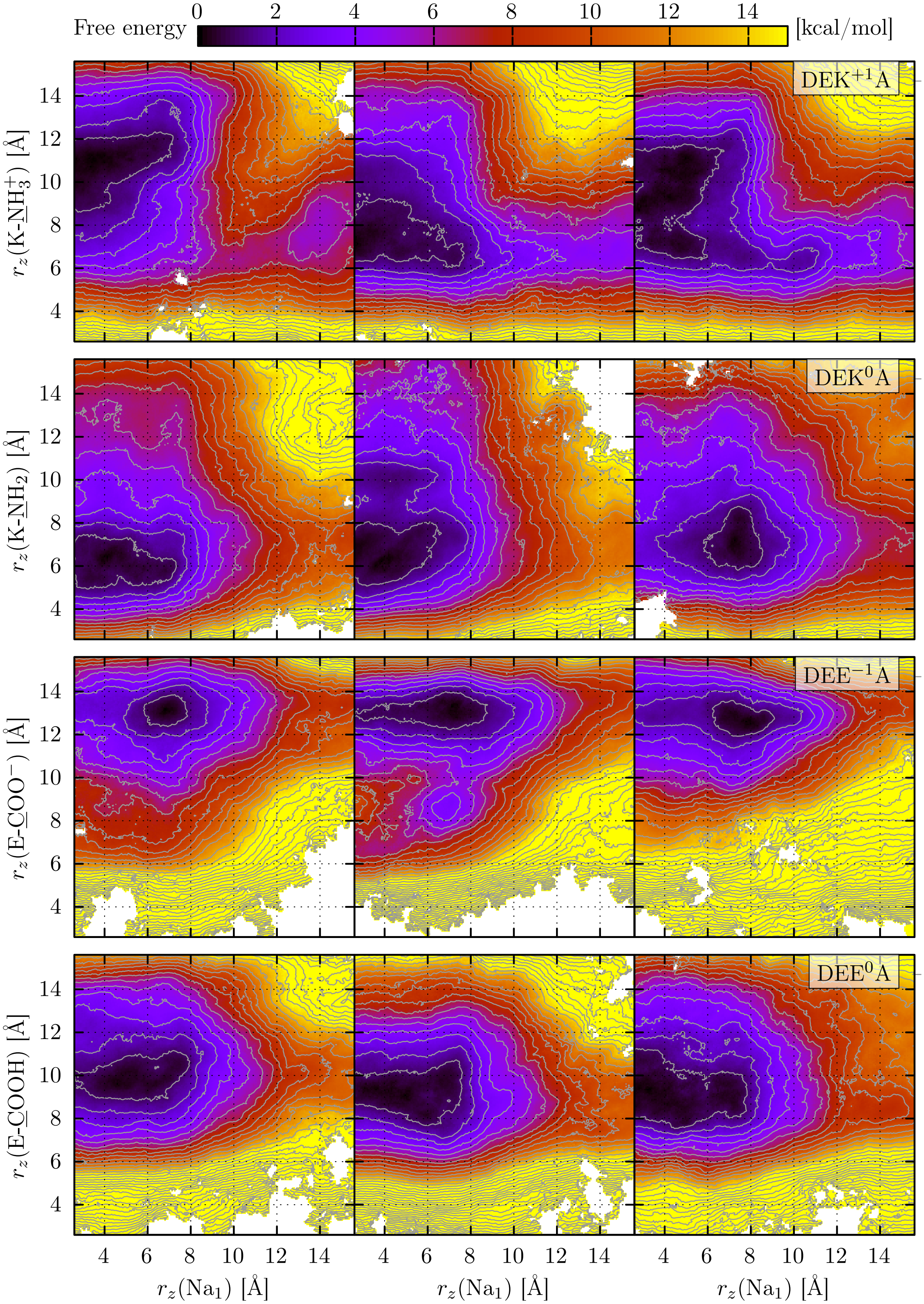


**Figure S6:** OTFP$_1$ FESs for single $Na^+$ translocation for each of the four systems of the Cryo-EM Na$_v$1.2 structure, as indicated in the labels.

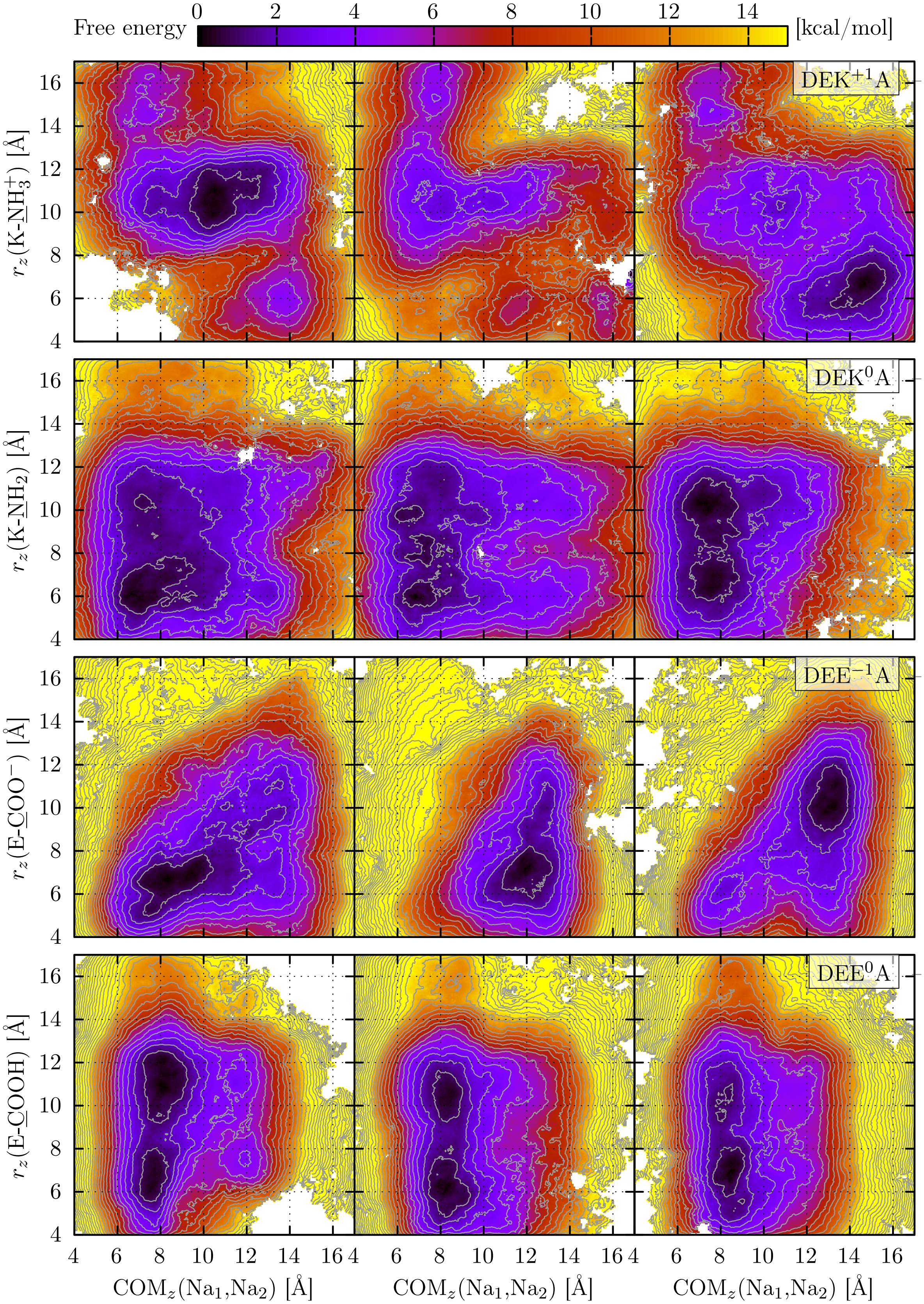


**Figure S7:** $OTFP_2$ FESs for single $Na^+$ translocation for each of the four systems of the $Na_v1.2$ model, as indicated in the labels.

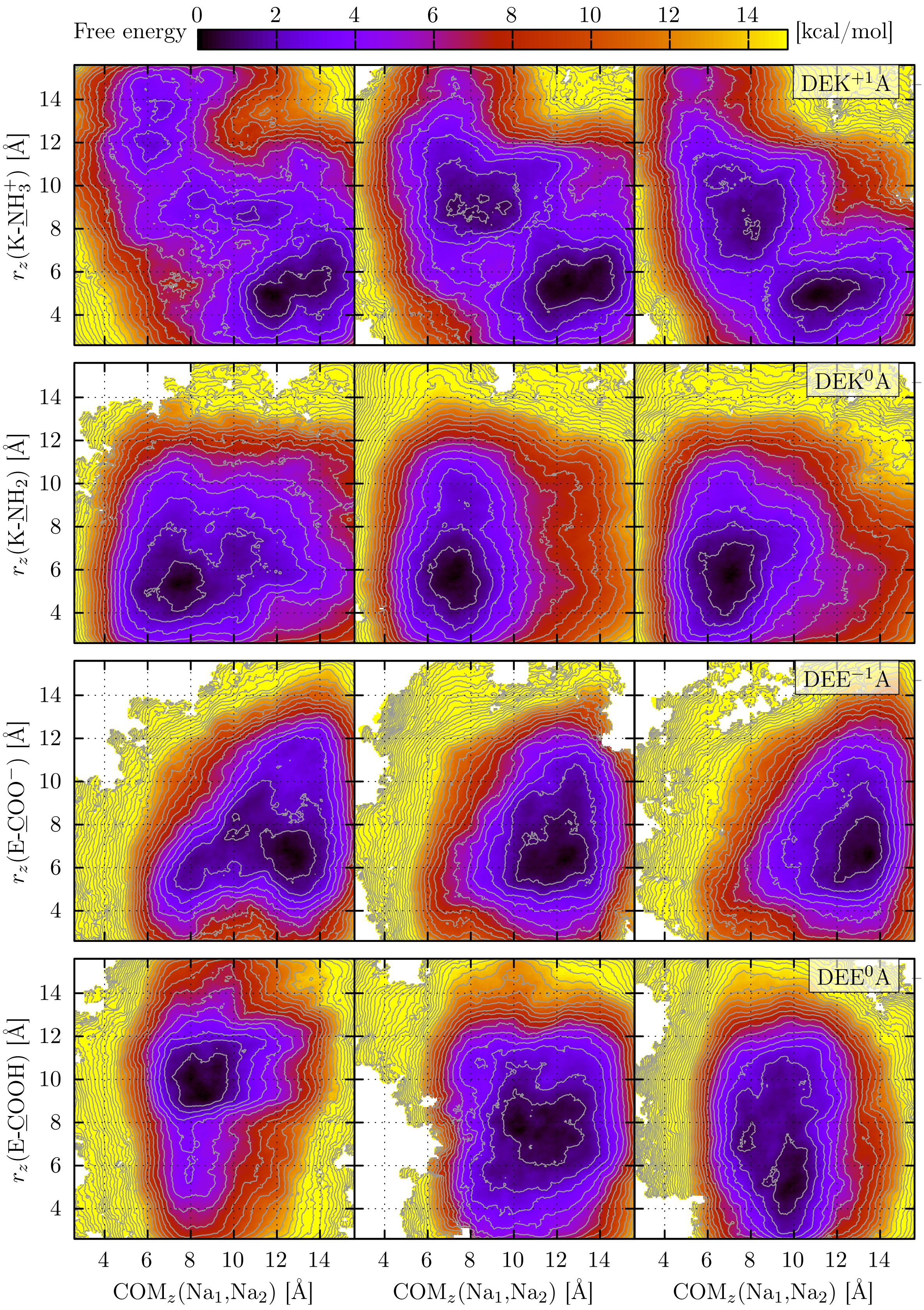


**Figure S8:** OTFP$_2$ FESs for single $Na^+$ translocation for each of the four systems of the Cryo-EM Na$_v$1.2 structure, as indicated in the labels.

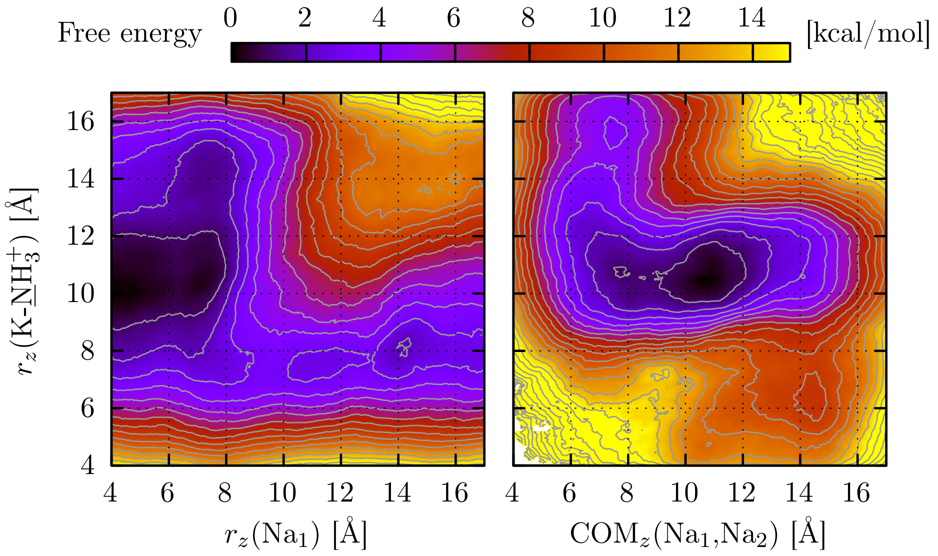


**Figure S9:** Average FESs based on the TAMD-OTFP simulations with a 150 mM NaCl concentration for the DEK$^+$A system based on the homology-based $Na_v1.2$ configuration ($OTFP_1$: left panel, $OTFP_2$: right panel). These simulations were performed with a larger rectilinear exclusion zone having corners at (-6, -6, -4) and (6, 6, 22) Å.

### List of Movies

Single ion permeation events observed for the various WT systems. In all the movies we represent the first three repeats, following the legend of **Figure 1**: domain I, gray; domain II, yellow; domain III, green. The fourth domain is excluded for clarity. The side-chains of the most external residues belonging to the EEDD motif are represented as red sticks. The charged residues of the three represented repeats belonging to the DEKA motif are also included (D and E are represented again with red side chains. The side-chain of the lysine is shown in blue sticks, with the NZ atom shown as a sphere). The internal carbonyl oxygen atoms of the two proceeding residues in each repeat are also included as red points. The permeating $Na^+$ ion is represented as an orange VdW sphere. Movies are available free of charge at `https://github.com/BeatriceCorradi/Nav_Channels_movies.git`

- **Movie S1**. Permeation of a single $Na^+$ ion in the MD simulation of the $Na_v$ 1.2 model (time-step $\Delta t = 2$ fs). Unrestrained production.
- **Movie S2**. Permeation of a single $Na^+$ ion in the MD simulation of the $Na_v$ 1.1 model (time-step $\Delta t = 4$ fs). Restrained equilibration.
- **Movie S3**. Permeation of a single $Na^+$ ion in the MD simulation of the $Na_v$1.2 model (time-step $\Delta t = 4$ fs). Restrained equilibration.
- **Movie S4**. Permeation of a single $Na^+$ ion in the MD simulation of the $Na_v$1.6 model (time-step $\Delta t = 4$ fs). Restrained equilibration.
- **Movie S5**. Permeation of a single $Na^+$ ion in the MD simulation of the $Na_v$1.2 Cryo-EM structure (time-step $\Delta t = 4$ fs - run 1). Unrestrained production.
- **Movie S6**. Permeation of a single $Na^+$ ion in the MD simulation of the $Na_v$1.2 Cryo-EM structure (time-step $\Delta t = 4$ fs - run 2). Restrained equilibration + unrestrained production.
